# Fear in action: Fear conditioning and alleviation through body movements

**DOI:** 10.1101/2022.06.20.496915

**Authors:** Maria Alemany-González, Martijn E. Wokke, Toshinori Chiba, Takuji Narumi, Naotsugu Kaneko, Hiraku Yokoyama, Katsumi Watanabe, Kimitaka Nakazawa, Hiroshi Imamizu, Ai Koizumi

## Abstract

Acquisition of fear memories enhances survival especially when the memories guide defensive movements to minimize harm. Accordingly, fear memories and body movements have tight relationships in animals: Fear memory acquisition results in adapting reactive defense movements, while training active defense movements to avoid threat reduces fear memory. However, evidence in humans is scarce because their movements are typically marginalized in experiments. Here, we tracked participants’ whole-body motions while they underwent fear conditioning in a virtual 3D space. First, representational similarity analysis of body motions revealed that participants obtained distinct spatiotemporal movement patterns through fear conditioning. Second, subsequent training to actively avoid threats with naturalistic defensive actions led to a long-term (24 hrs) reduction of physiological and embodied conditioned responses, while extinction or vicarious training only transiently reduced the responses followed by their spontaneous return. Together, our results highlight the intrinsic role of body movements in human fear memory functions, suggesting the potential for improving fear memory interventions through embodiment.

## Introduction

Threatening experience induces learning of threat-predictive cues in the environment. This learning capacity - threat conditioning - enhances the chance of survival by minimizing direct contact with the threats (*1*–*3*). In the real world, minimization of harm requires bodily defensive movements to act on external situations such as attacking or escaping from threats (*4*, *5*) with rich movement repertoires (*6*–*10*). Accordingly, threat conditioning in animals results in the emergence of defensive body movements such as freezing (*11*–*13*). This highlights that optimization of defensive actions to better cope with threats serves the ultimate goal of threat conditioning.

Interestingly, training defensive movement *per se* results in the alleviation of threat-conditioned responses in animals (*2*). For instance, conditioned freezing responses in rats are reduced by training active-avoidance behaviors, such as escaping to terminate a threat-conditioned cue and prevent threatening electric shock (*14*, *15*). Learning to alternate from passive to active defensive responses is an intrinsic capacity of the brain where a prefrontal top-down mechanism facilitates neural pathways prioritizing the latter over the former (*2*, *9*, *16*).

Unlike the demonstrated roles of body movements in both acquisition and alleviation of threat-associative memories in animals, evidence in humans is scarce because their body movements are often marginalized in research (*17*, *18*). First, acquisition of threat conditioning in humans is typically assessed with peripheral physiological responses such as skin conductance responses (*19*) and heart rates (*20*, *21*). Although those physiological responses may emerge as preparatory modulators of defensive bodily movements (*22*–*24*), they imply little about overt bodily movements.

Second, although the active-avoidance procedure in animals (*14*, *15*) has been translated to humans in seminal works (*16*, *25*–*29*), procedures in humans are generally less embodied than in animals. For example, participants avoid aversive shocks through finger key-pressing to control icons on a monitor that are unrelated to any ecologically-valid defensive action (*25*, *29*). Those procedures successfully alleviated fear (*16*) with a long-term effect (24 hrs) (*25*, *28*–*30*), which outperformed the effects of extinction procedures to simply repeat conditioned cues without further reinforcement with threats (*25*). Nevertheless, examining the efficacy of embodied active-avoidance procedures in alleviating fear memory is a critical step forward in their clinical applications. Because post-traumatic psychopathologies emerge from various threatful experiences that require bodily actions for avoidance - such as interpersonal violence, looming vehicles, and natural disasters (*31*).

The scarcity of body movement measures in studying human emotions (*18*, *32*) partly owes to a common technical limitation. That is, experimental control is typically achieved by constraining participants in a seated or laid-down posture. This limits the available actions to peripheral movements such as fingers and gaze, hindering the measurements of threat-conditioned movements and the implementations of bodily defensive actions.

To overcome these limitations, we developed a spatiotemporally dynamic threat conditioning paradigm with virtual reality (VR) (*18*, *33*–*36*) to unleash participants’ naturalistic body movements. Moreover, we monitored participants’ whole body movements with a motion-tracking system. For threat conditioning, participants experienced a simulated real-life traumatic scenario of interpersonal violence where a specific avatar (CS+) inflicts threatening violence (US; unconditioned stimulus). Participants then underwent training to avoid violence through naturalistic bodily defensive movements using a virtual security spray against the CS+ avatar (**Fig 1**). We chose a scenario of interpersonal violence because it is one of the most common traumatic events in modern societies e.g., affecting 21-80% of adolescents (*37*, *38*) which can lead to post-traumatic stress disorders (PTSD) (*39*).

**Figure 1.**
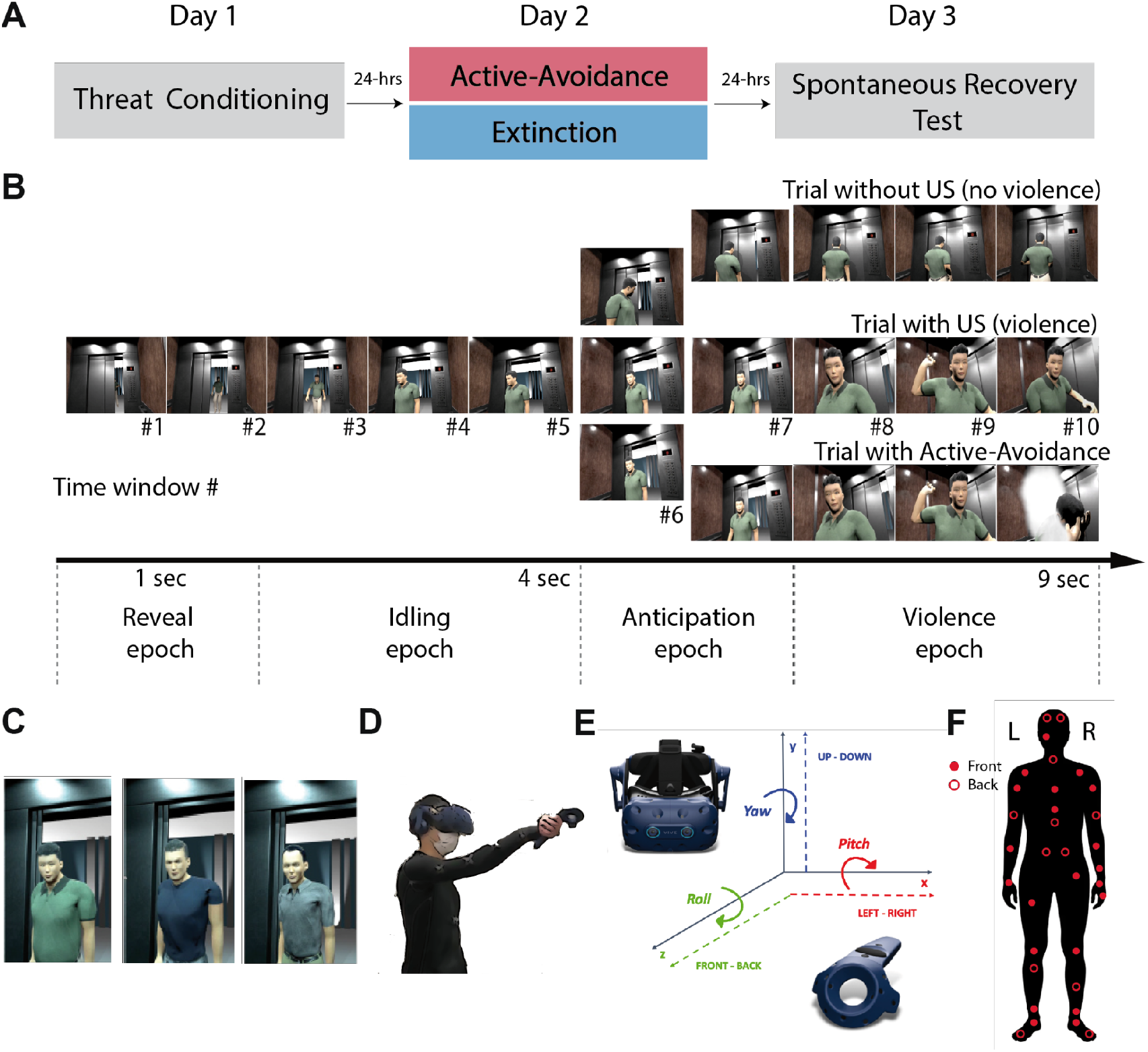
Schematics of the experimental design. **(A)** Experimental sessions across three consecutive days. **(B)** Virtual scenes for non-violent trials (upper), violent trials (middle), and embodied active-avoidance trials (lower) progressed across 10 representative time windows within a trial (see Movies S1-3). **(C)** Three male avatars, two of which served as CS+s and another as CS- in a counterbalanced manner across participants. **(D)** Illustration of a subject performing a defense movement in the embodied active-avoidance training (see Movie S2). **(E)** Schematics of the three-dimensional spaces capture the participant’s body movements as tracked in the head-mount display and a hand-held controller. **(F)** Marker distribution on the full-body motion tracking system.

Anticipating the results, participants obtained distinct multidimensional body movement patterns as threat-conditioning progressed, suggesting the acquisition of embodied conditioned responses. Furthermore, training embodied active-avoidance actions led to a long-term (24 hr) reduction of both physiological and embodied conditioned responses while control extinction or vicarious training led to only a transient reduction of the responses followed by their spontaneous return. The study suggests how reviving the roles of bodily actions could benefit the understanding and treatment of human traumatic memories.

## Results

### Experimental design

We conducted experiments across three consecutive days (**Fig 1A**). On day 1, 41 participants (Group 1) underwent a naturalistic threat conditioning session in an unconstrained standing posture inside a virtual elevator. The body movements of participants were constantly reflected in the movements of their avatar-self, whose appearance was semi-customized to reflect their gender (see Mirror Sessions in Methods).

Two male avatars (CS+s), but not a third male avatar (CS-), intimidated and hit the participants virtually on proportions of the trials (**Fig 1B and C**, see **Movie S1**). Here, violence by CS+ avatars served as an unconditioned stimulus (US). On day 2, participants underwent embodied active-avoidance training (ACT training) where they could defend themselves with a security spray to prevent violence (US) from one of the CS+ avatars (CS+_active_) (**Movie S2**). Specifically, the hand-held controller in their right hand appeared as a virtual security spray. Upon seeing the precursive action for violence by the CS+_active_ avatar (i.e., raising his hand), participants positioned their bodies and pulled the lever on the controller to virtually emit the white spray against the avatar to prevent violence (**Fig 1D**). In conventional extinction training (EXT training), participants were repetitively exposed to the other CS+ (CS+_extinction_) avatar without further reinforcement by violence (**Movie S3**). On day 3, participants encountered both CS+_active_ and CS+_extinction_ avatars without violence and we tested for the return of threat-conditioned responses in the spontaneous recovery test session. Along with physiological defensive reactions as measured with skin conductance responses (SCR) and subjective ratings of fear and violence-likelihood, the body movements of the participants were tracked with the VR head-mount-display and hand-held controller (**Fig 1E**) or with a full-body motion system (**Fig 1F**) (see Methods). All sessions included trials without US and only those trials were analyzed to assess the anticipatory conditioned responses.

### Threat conditioning in subjective and physiological measures

With VR threat-conditioning, participants successfully acquired conditioned responses (CR; larger responses with CS+ than CS-), as evidenced by their subjective ratings of fear (**Fig 2A**). After the conditioning session, participants reported higher feelings of fear as well as higher anticipation of US (violence) with both CS+_active_ and CS+_extinction_ relative to CS- [Fear: CS+_active_ *t*(40) = 6.44 *p* < 0.001; BF_+0_ >100; CS+_extinction_ *t*(40) = 8.00 *p* < 0.001; BF_+0_ >100; US likelihood: CS+_active_ *t*(40) = 9.97, *p* < 0.001; BF_+0_ >100; CS+_extinction_ *t*(40) = 11.72, *p* < 0.001; BF_+0_ >100; all one-tailed for predicted positive CR]. Subjective ratings were statistically similar between CS+s [CS+_extinction_ Vs CS+_active_; Fear: *t*(40) = -0.601, *p* = 0.55; BF_+0_ = 5.003; US-likelihood: *t*(40) = 0.07, *p* = 0.93; BF_+0_ = 5.91].

**Figure 2.**
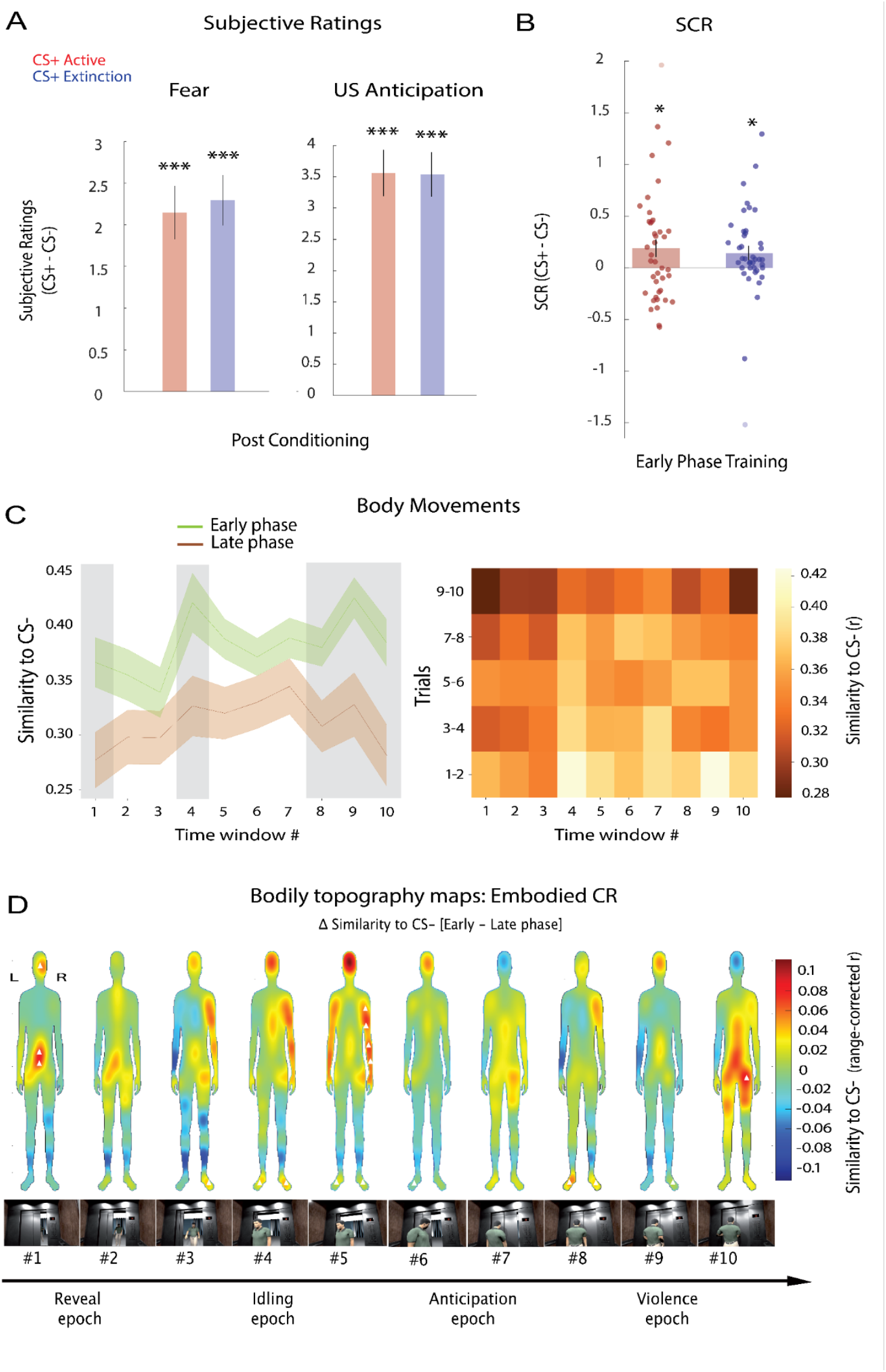
Threat conditioning in subjective ratings, SCR, and body movements. **(A)** Conditioned responses [CR: CS+ versus CS-] in subjective ratings of fear and US-anticipation on day 1. Values are baseline-corrected with pre-conditioning ratings. **(B)** CR in SCR during the early phase (trials 1-2) of the training on day 2. **(C)** Embodied CRs captured as a decrease of similarity in body movement patterns toward the CS+s versus the CS- avatars from the early (trials 1-2) to late phase (trials 9-10) (left) or across all phases of the threat conditioning session (right). The shaded areas in the left panel indicate embodied CRs with moderate and strong evidence (BF > 3). **(D)** Maps depicting embodied CRs as threat conditioning progressed from the early to late phase. In maps, the position of each bodily ROI was approximated with one of its constituent markers. Warmer colors indicate larger embodied CRs. For the demonstrative purpose, values (similarity to CS-, r) were corrected to restore the original min-max range after spatial smoothing per time window. Triangles (▴) in the maps indicate the ROIs with anecdotal or moderate evidence based on BF. ROIs with only positive conditioning effects are shown with one-tailed Bayesian analysis for interpretation. For both SCR and body movements analysis, only the trials without violence (US) were analyzed. Data are represented as mean ± between-subjects SEM. \**p* < 0.05 \*\*\**p* < 0.001.

Participants also displayed physiological CR in SCR measures 24 hrs following the conditioning session (**Fig 2B**). Specifically, CRs to both CS+_active_ and CS+_extinction_ were significant during the early phase of the training sessions on day 2 [CS+_active_ *t*(40) = 2.23 *p* = 0.016; BF_+0_ = 3.05; CS+_extinction_ *t*(40) = 2.05 *p* = 0.02; BF_+0_ = 2.18; all one-tailed], with no statistical difference between the two CS+s [*t*(40) = 0.35 *p* = 0.72; BF_01_ = 5.58]. During the conditioning session on day 1, SCR with both CS+_active_ (M = 0.24 ± s.e. 0.02) and CS+_extinction_ (0.24 ± 0.02) were higher than CS- (0.23 ± 0.02) on trials without US, indicating numerical yet non-significant CRs [CS+_active_ *t*(40) = 0.91, *p* = 0.18; BF_+0_ = 0.403; CS+_extinction_ *t*(40) = 0.71, *p* = 0.23; BF_+0_ = 0.32; all one-tailed] with no statistical difference between the two CS+s [*t*(40) = 0.13 *p* = 0.89; BF_01_ = 5.88] (**Fig S1A**). This overnight enhancement of CR from day 1 to day 2 likely reflects fear memory consolidation (*40*–*43*). Our subsequent analyses were inclusive for all participants because those with less differential CR could be more prone to anxiety and thus represent a relevant cohort in fear memory research (*20*, *44*).

### Threat conditioning of human body movements

Next, we asked whether participants also developed threat-conditioned body movement patterns. Using representational similarity analysis (RSA) (*45*–*48*), we examined whether the body movement similarity between the encounterance with CS+ versus CS- avatars decreased from the early to late phase of the conditioning session. A decrease in the similarity indicates the acquisition of distinctive body movement patterns for CS+ relative to CS-, representing embodied CRs. As in the analyses with SCR, to assess the anticipatory conditioned responses, only the trials without US were analyzed where both CS+ and CS- avatars displayed identical motion trajectories.

For RSA, body movement patterns were first quantified with the position and rotation on the 3D axes of the head and right hand (12 dimensions), which were tracked with the VR head-mount-display and hand-held controller (**Fig 1E**). To obtain a measure that is independent of coordinate values and unbiased by previous positions during a trial, we then calculated the absolute difference between two consecutive recorded samples (|Position(t2)-Position(t1)|) within trials, reflecting the magnitudes of movement per sample frame. To reduce the number of comparisons while retaining enough temporal resolution to examine unique epochs of the scene (e.g. anticipation of violence), the spatial pattern similarity was evaluated for 10 unique time windows within each trial (9 sec in total); These captured *reveal (*Time windows *#1,2), idling (#3-5), anticipation (#6,7), and violence epochs (#8-10)* (**Fig 1B**, see **Movies S1-3**).

The embodied CRs were captured with strong evidence in four threat-relevant time windows (**Fig 2C**), as indicated by Bayes Factors (BF). These included the period of the initial encounter with the CS avatars (*reveal-epoch*) and the last time windows, which correspond to the period of potential violence occurrence (*violence-epoch*; time windows #1, 8, 9, & 10: *t*-values(40) > 3.26, *p*-values < 0.001 FDR-corrected < 0.05, Cohen’s d > 0.51; all BF_+0_ > 29.46, all one-tailed given predicted embodied CRs with decreasing similarity, see **Fig 2C** for the results in other time windows). For time window #4, we observed a moderate effect [BF_+0_ = 5.85] which did not survive the multiple comparison corrections with frequentist statistics [*t*(40) = 2.56, *p* = 0.007 FDR corrected > 0.05, Cohen’s d = 0.39]. These findings show that movement patterns in response to CS+ and CS- became increasingly dissimilar when fear conditioning was established.

The results were qualitatively similar between the two CS+ avatars examined separately (**Fig S1B**). Embodied CRs with CS+_active_ were found with strong evidence in time windows #1 & 10 [*t*-values(40)> 2.97, *p*-values < 0.002 FDR corrected < 0.05, Cohen’s d > 0.464; BF_+0_ > 14.92; all one-tailed] and with moderate evidence in time windows #4 & 8 [*t*-values(40)> 2.70, *p*-values < 0.005 FDR corrected < 0.05, Cohen’s d > 0.422; BF_+0_ > 8.05; all one-tailed]. All other times windows had BF_+0_ < 1.11. Similarly, embodied CRs with CS+_extinction_ were strong in time windows #1, 9 & 10 [*t*-values(40) > 3.28, *p*-values < 0.001 FDR corrected < 0.05, Cohen’s d > 0.513; BF_+0_ > 31.25; all one-tailed] and moderate in time windows #3, 5 & 8 [*t*-values(40)> 2.58, *p*-values<0.007 FDR corrected>0.05, Cohen’s d >0.403; BF_+0_>6.16; all one-tailed]. All other time windows had BF_+0_ < 1.20 (**Fig S1B**).

The results were specific to threat conditioning effects as they were not replicated when examining the deviation of movement patterns between the two CS+s. The changes in the similarity between CS+_active_ and CS+_extinction_ from the early to late phase of the conditioning session were found only in the less threat-relevant *reveal-* and *idling-epochs* [time windows #1 & 5: *t*-values(40) > 3.32, *p*-values < 0.002 FDR corrected < 0.05, Cohen’s d > 0.51; BF_10_ > 17.12]. Importantly, the similarity between the two CS+s encounters did not decrease in later time windows when participants anticipate potential violence in *anticipation* and *violence-epochs* [time windows #6-10: *t*-values(40) < 2.23, *p*-values > 0.032 FDR corrected > 0.05, Cohen’s d < 0.35; BF_10_ < 1.53]. While a generic deviation of body movement patterns emerges when an avatar’s identity is revealed with the elevator door opening, embodied CR may emerge specifically in time windows when potential threats are anticipated.

To unpack the spatial specificity of embodied CRs, we ran RSAs for the head and right hand separately (**Fig S1C**). The results suggested that embodied CRs were more pronounced in the hand than in the head. For hand movement, we found moderate to strong embodied CRs in time windows including the threat-relevant *violence-epoch* #4, 5, 8 & 9 [*t*-values(40)> 2.75, *p*-values<0.004 FDR corrected<0.05, Cohen’s d > 0.43; BF_+0_>8.92, all one-tailed]. For head movement, however, we found moderate to strong CRs only in time windows less relevant to threat #1 & 10 [*t*-values(40) > 2.59, *p*-values < 0.007 FDR corrected < 0.05, Cohen’s d > 0.40; BF_+0_ > 6.23, all one-tailed].

To further elucidate the spatio-temporal dynamics of embodied CRs across the entire body, a second group of 53 participants (Group 2) completed the same conditioning session as Group 1 with an addition of a whole-body motion tracking system (**Fig 1F**) (see *Movement Representational Similarity Analysis* in Methods). We conducted searchlight RSAs with 31 bodily regions-of-interests (ROIs), each composed of the nearest three body markers tracking their 3D positions. The bodily topographical maps, inspired by a previous study (*49*), capture the gradients of CRs across the whole body and time windows (**Fig 2D**). While the maps serve demonstration purposes, we calculated Bayesian factors for each ROI and time window. Of note, similarly to Group 1, moderate embodied CRs were captured in the right arm toward the end of *idling-epoch* [time window #5: 1 ROI BF_+0_=3.64; one-tailed]. This CR was spatially distributed over the contiguous ROIs in the right arms with anecdotal evidence [time window #5: 3 ROIs BF_+0_>1.24; one-tailed]. We additionally captured moderate CRs in bilateral feet in windows including *idling and violence-epochs* [time windows #4 & 8: 2 ROIs BF_+0_>3.22; one-tailed] and anecdotal CRs in contiguous ROIs in the feet [time window #3-6, 8 & 9: 2 ROIs BF_+0_>1.40; one-tailed] (**Fig 2D**) (see Supplementary for the full descriptions of statistical results).

Together, threat-conditioning altered spatiotemporal dynamics of body movements, signaling embodied threat-anticipatory responses in human participants.

### Fear reduction effects through embodied active-avoidance

Next, we asked if the CRs can be alleviated through the embodied procedure to actively avoid harm (violence) inflicted by the CS+ avatar. For this, participants physically defended against one of the two CS+s avatars (CS+_active_) in the embodied active-avoidance training (ACT training) while they merely observed the other CS+ avatar (CS+_extinction_) without being reinforced by violence in the conventional extinction training (EXT training) on day 2 (**Fig 1B**). The long-term (24 hrs) effects of training were tested in the early phase of the spontaneous recovery test on day 3 (**Fig 1A**) because the spontaneous return of CRs typically emerge only in the first few trials and quickly fade through additional extinction in subsequent unreinforced trials (*50*, *51*).

Critically, we found that the long-term effect of reducing embodied CRs was more pronounced with ACT than with EXT training. In the early phase of the test, body movement patterns during CS+ encounters became more similar to CS- encounters after ACT training than EXT training (**Fig 3A**). This long-term effect to neutralize the body movements was observed within the threat-relevant *anticipation-epoch,* as indicated by moderate evidence [time windows #6 & 7: *t*(40) = 2.53, *p* = 0.008 FDR corrected >0.05, Cohen’s d = 0.40, BF_+0_ = 5.46; *t*(40) = 2.26, *p* = 0.015 FDR corrected >0.05, Cohen’s d = 0.35, BF_+0_ = 3.20; for all other times windows BF_+0_ < 1.01; all one-tailed for predicted advantage of ACT training (*25*).

**Figure 3.**
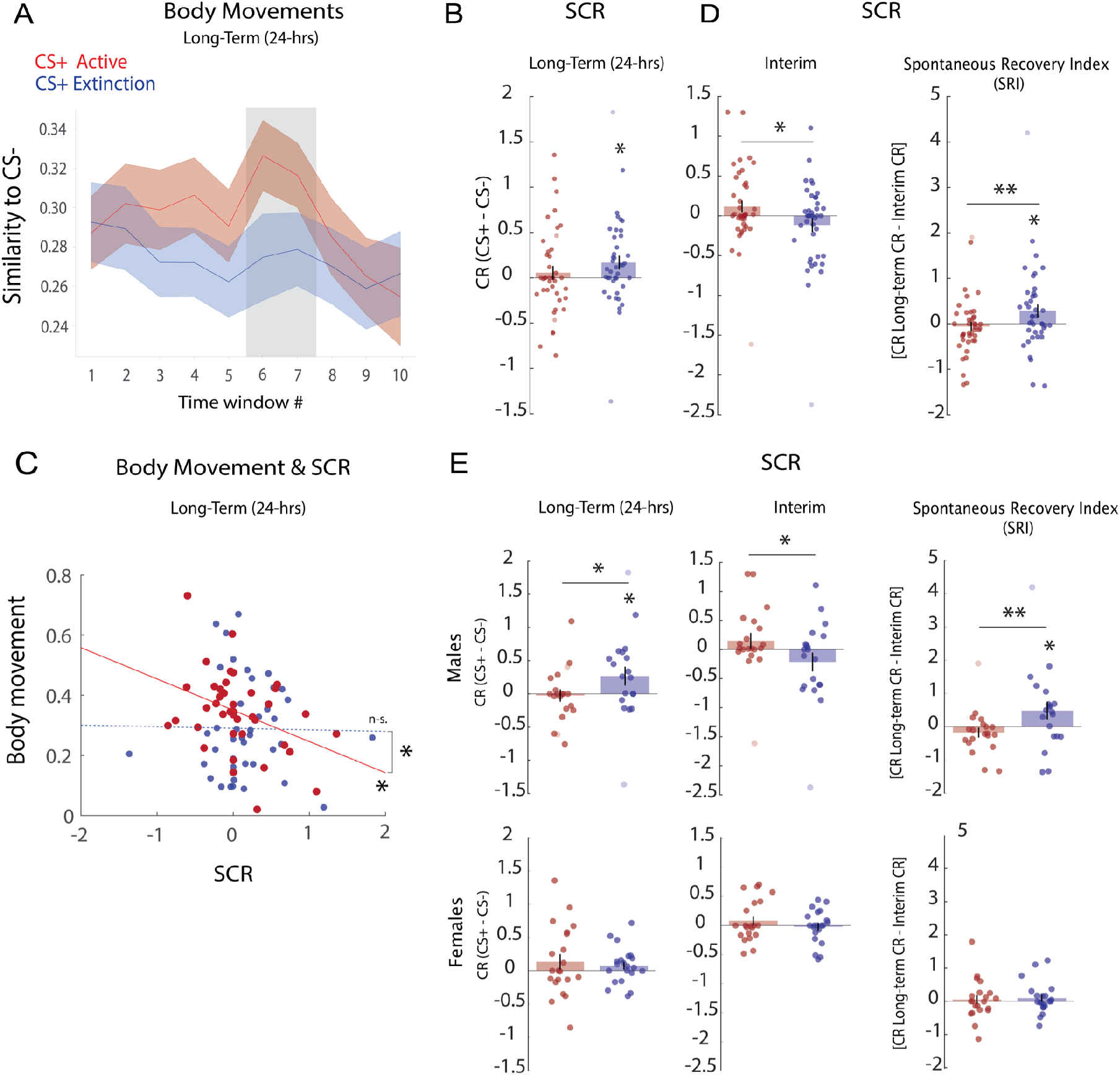
Comparison of fear alleviation effects between embodied active-avoidance (ACT) training and Extinction (EXT) training. **(A)** Long-term effects of ACT and EXT training on body movement patterns. Movement patterns during CS+s encounters became more similar to CS- encounters after the ACT training compared to the EXT training in *anticipation-epoch* (windows #6 & 7) as indicated by moderate evidence with Bayes factors. **(B)** Long-term effects of ACT and EXT training on conditioned SCR. **(C)** Correlation between CR in SCR and body movements in the Spontaneous Recovery Test 24 hrs after the training. Scatter plots with least-square regression lines of the intra-subject correlations between SCRs and body movements. **(D)** Interim effects of ACT and EXT training on conditioned SCR (left panel) and spontaneous recovery index (SRI) reflecting the difference between the interim versus the long-term conditioned SCR (right panel). **(E)** The ACT and EXT training Interim and long-term effects and on SRI in female and male gender groups. All panels in Fig 3 include only the trials without US. Error bars represent the SEM. Dots superimposed on bar graphs represent individual participants’ data where outliers were included in the analyses yet faded in their colors for the demonstrative purposes. Data are represented as mean ± between-subjects SEM. \**p* < 0.05 \*\**p* < 0.01. The shaded areas in (A) indicate moderate and strong evidence (BF > 3).

Similarly to the body movement results, the long-term reduction of SCR was more pronounced with ACT training than EXT training (**Fig 3B**). In the early phase of the test, only CS+_extinction_ induced a significant CR [*t*(40) = 2.21, *p* = 0.016; BF_+0_ = 2.21] while CS+_active_ did not [*t*(40) = 0.76, *p* = 0.22; BF_+0_ = 0.34; all one-tailed]. Interestingly, the long-term effect of ACT training on SCR significantly correlated with the effect on embodied CRs (**Fig 3C**). In the spontaneous recovery test, participants with weaker CR to CS+_active_ also displayed more neutralized body movement patterns to CS+_active_ (i.e., increased similarity to CS-) [r = -0.37, *p* = 0.015]. This correlation was absent with CS+_extinction_ [r = -0.01, *p* = 0.92] with a significant difference between the CS+s [r_active_ Vs r_extinction_ z = -1.65, *p* = 0.047]. The embodied ACT training may rely on unique fear reduction mechanisms that can neutralize conditioned body movements along with the reduction of physiological CR.

We next asked if this advantage of ACT training already emerged during the training sessions on day 2. For this, we examined SCR in the late training phase to assess the interim effects of each training (**Fig 3D left**). We did not assess the interim effects on embodied CRs because the task instruction for specific defensive actions itself altered the body movement patterns, hindering access to the embodied CRs during the training (see **Fig S2**). Critically, with SCR, CRs to both CS+_extinction_ and CS+_active_ were no longer significant in the late training phase [CS+_extinction_ *t*(40) = - 1.35, *p* = 0.90; BF_+0_ = 0.07; CS+_active_ *t*(40) = 1.52, *p* = 0.06; BF_+0_ = 0.90; all one-tailed], suggesting that both ACT and EXT training achieved interim effects in reducing SCR. However, CR to CS+_extinction_ was lower than CS+_active_ [*t*(40) = 2.408, *p* = 0.02; BF_10_ = 4.31] (**Fig 3D left**). This suggests that although ACT training resulted in a better long-term reduction of SCR on day 3, EXT training was transiently more advantageous in reducing SCR to CS+_extinction_ on day 2. This may not be surprising because the CS+_active_ avatar continued to display precursive actions for violence (US) during the training on day 2.

The ACT training effect may outperform the EXT training effect overnight because extinction-based procedures are prone to spontaneous recovery of once-extinguished fear responses (*52*). To test this possibility, we quantified the increase of CRs from the late training on day 2 (interim effect) to the early spontaneous recovery test on day 3 (long-term effect) as spontaneous recovery index (SRI) (*25*). In line with previous studies (*25*, *52*), SRI was significant with CS+_extinction_ [*t*(40) = - 1.97, *p* = 0.02; BF_+0_ = 1.90; one-tailed]. In contrast, SRI was not significant with CS+_active_ [*t*(40) = 0.59, *p* = 0.72; BF_+0_ = 0.11; one-tailed]. Critically, SRI with CS+_active_ was significantly smaller than CS+_extinction_ [*t*(40) = -2.57, *p* = 0.007; BF_+0_ = 6.13; one-tailed] (**Fig 3D right**).

Together, although EXT training transiently led to a better interim fear reduction effect on day 2 than ACT training, this advantage of EXT training was transient as its effect was susceptible to spontaneous recovery 24 hrs later. Meanwhile, although the interim effect of ACT training was transiently weaker than EXT training, the fear reduction effect of ACT training was more stable over 24 hrs to outperform the long-term effect of EXT training by resisting spontaneous recovery. This long-term advantage of ACT training was robust to the training order (**Fig S3A**) and outliers (**Fig S3B**) (see **Fig S4** for training effects on subjective ratings).

The ACT training may resist spontaneous recovery of CRs because it relies on a unique fear reduction mechanism than the EXT training. Typically, a conventional extinction procedure is prone to the return of extinguished fear responses because it relies on the mechanisms to suppress original fear memories (*53*). In line with this, participants who showed smaller CRs to CS+_extinction_ during EXT training on day 2 showed a larger return of CRs on day 3 [R = -0.58, *p* < 0.001; BF_+0_ = 529.8] (**Fig S3C**). This suggests that stronger suppression of CRs through EXT training may result in a stronger return of CRs 24 hrs later. Critically, such a correlation was absent with ACT training [r = 0.12, *p* = 0.42; BF_+0_ = 0.26] with a significant difference between the training types [r ACT Vs r EXT z = -3.41, *p* = 0.002], suggesting that ACT training may be more independent from the suppressive mechanisms.

Finally, given the gender differences in treatment responses among patients with PTSD (*54*), we examined whether the training effects differed between genders with a conventional physiological measure of SCR from Group 1 (21 females, 20 males). This revealed a significant interaction between gender and training in the long-term effect (**Fig 3E**) [*F*(1,39) = 4.76, *p* = 0.03]. Only males showed a significantly better long-term reduction of CR through ACT than EXT training [*t*(19) = - 2.51, *p* = 0.02; BF_+0_ = 2.77] when females did not [*t*(20) = 0.53, *p* = 0.601; BF_+0_ = 0.25] (**Fig 3E left**). The analysis of SRI also revealed a significant interaction between gender and training type [*F*(1,39) = 5.94, *p* = 0.01] (**Fig 3E right**), suggesting how the interim effect of EXT training was outperformed by ACT training in long-term (24 hrs), especially among males. Only with males, SRI was significantly higher with CS+_extinction_ than CS+_active_ [*t*(19) = -3.27, *p* = 0.004; BF_+0_ = 11.18; *t*(19) = -3.61, *p tukey* = 0.004] while not with females [*t*(19) = -0.25, *p tukey*= 0.99]. Gender did not interact with the interim effects alone (**Fig 3E middle**). These results suggest that the embodied ACT training may potentially benefit males more than females.

### Fear reduction effects depend on embodiment of active-avoidance

Do fear reduction effects of ACT training depend on whether participants defend against threats through their own body movements? To test this, participants in Group 2 with full-body motion tracking (**Fig 1F**) underwent the ACT training either with or without their own movements in two subgroups (**Fig 4A**): Self-group physically defended themselves with a virtual security spray to prevent violence as in Group 1 **(Fig 1D),** whereas Vicarious-group merely observed a third avatar defending with a security spray on their behalf (**Fig 4B)**. Both groups underwent the ACT training with one of the two CS+s (CS+_active_) while they underwent no training with the other CS+ (CS+_untrained_). The training effect was compared between the two subgroups in the spontaneous recovery test 24 hrs later.

**Figure 4.**
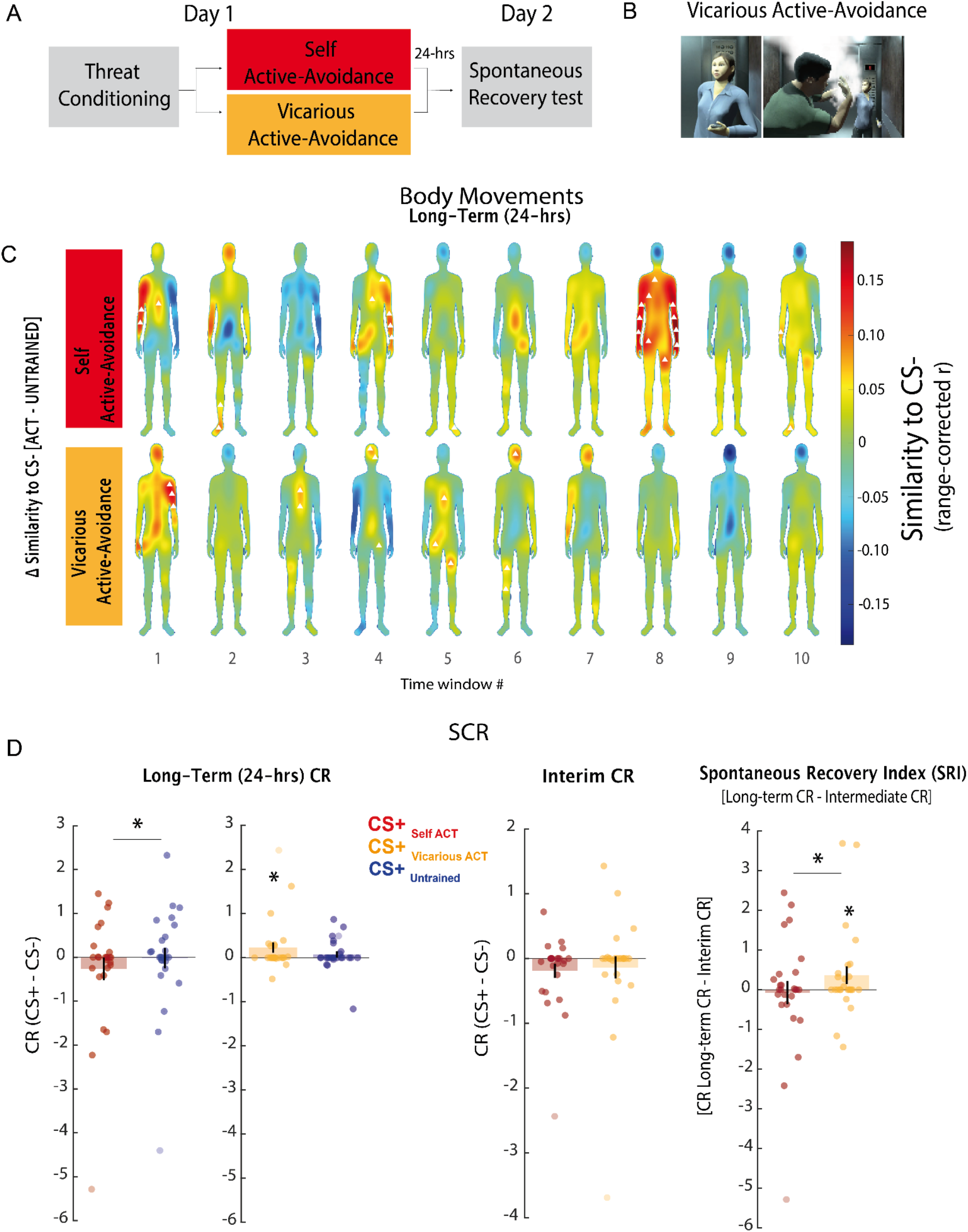
Comparison of fear alleviation effects between Self and Vicarious ACT training. **(A)** The flow of the experiment for the second group across two consecutive days. **(B)** Female passenger avatar defending against the violent CS+_active_ avatar with a security spray on the participant’s behalf in the Vicarious training. **(C)** Maps depicting ACT training effects by comparing body movements with CS+_active_ and CS+_untrained_ in the spontaneous recovery test session for Self- (upper) and Vicarious-group (lower). Warm colors indicate that the ACT training rendered body movements with CS+ encounters more similar to CS- encounters compared to no training. Triangles (▴) in the maps indicate the ROIs with anecdotal, moderate, or strong evidence based on BF, where only ROIs with increased similarity to CS- movements compared to no training are shown for interpretation. **(D)** Interim and long-term effects of Self and Vicarious training on conditioned SCR (left and middle) and spontaneous recovery index (SRI) (right). All panels in Fig 4 include only the trials without US. Data are represented as mean ± between-subjects SEM. Dots superimposed on bar graphs represent individual participants’ data where outliers were included in the analyses yet faded in their colors for the demonstrative purposes. \**p* < 0.05

Critically, we found the superiority of the Self-group relative to the Vicarious-group in neutralizing the conditioned body movements. The bodily topographic maps (**Fig 4C**) demonstrate the effects in each group. Specifically, in the spontaneous recovery test, Self-group showed training effects to neutralize body movements (i.e., increase in movement similarity to CS- relative to no-training) in several ROIs in bilateral arms during *violence-epoch* as indicated by moderate evidence [time window #8: 6 ROIs BF_+0_ > 3.10; one-tailed] and by anecdotal evidence [3 ROIs BF>1.25]. Additionally, Self-group showed effects in *reveal-epoch* in the left arm and feet as indicated by moderate and strong evidence, respectively [Left arm: time window #1: 1 ROI BF_+0_=5.12; Left feet: #2: 1 ROI BF_+0_=63.25; all one-tailed] (see Supplementary for further results).

On the contrary, Vicarious-group did not show any apparent training effect in any body part during *violence-epoch*. A single moderate training effect was observed only in the right leg in *idling-epoch* [time window #5: 1 ROI BF_+0_=4.66], with additional anecdotal effects in a few spatially sparse ROIs during *anticipation-epoch* (see Supplementary). Together, the results suggest that the embodiment of ACT training in Self-group contributed to the long-term neutralization of body movement patterns.

Moreover, long-term reduction of SCR was more robust in Self-than in Vicarious-groups (**Fig 4D left**). In the early phase of the spontaneous recovery test, only Vicarious-group showed a significant CR to CS+_active_ when Self-group did not [Vicarious: *t*(25) = 1.96, *p* = 0.03; BF_+0_ = 2.08; Self: *t*(25) = -1.02, *p* = 0.84; BF_+0_ = 0.11; all one-tailed]. Moreover, only Self training significantly reduced CR toward CS+_active_ relative to CS+_untrained_ [*t*(25) = -1.91, *p* = 0.03; BF_10_ = 1.91] when Vicarious training did not [*t*(25) = 1.25, *p* = 0.22; BF_10_ = 0.47] (**Fig 4D left**). The interaction between group and CS+ type was significant for the long-term effect [*F*(1,50) = 5.03 *p* = 0.02]. Accordingly, SRI was only significant in the Vicarious-group [*t*(25) = -2.36, *p* = 0.02; BF_+0_ = 3.90; one-tailed] while not significant in the Self-group [*t*(25) = 1.42, *p* = 0.16; BF_+0_ = 0.21; one-tailed]. The long-term advantage of Self-group was also captured by a lower SRI to CS+_active_ (return of CR from day 2 to day 3) than Vicarious-group [*t*(52) = -2.20, *p* = 0.03; BF_10_ = 1.97] (**Fig 4D right**), with a significant interaction between group and CS+ type [*F*(1,50) = 4.68 *p* = 0.03] (see Supplementary for further results and **Fig S6** and **S7** for additional comparisons between the groups on SCR and subjective ratings). Together, these results suggest that long-term fear reduction effect was specific to the embodied active-avoidance training, as the less-embodied control training with vicarious defense training (Group 2) as well as extinction training (Group 1) only led to a transient reduction of fear which returned 24 hrs later.

## Discussion

By liberating and tracking naturalistic body movements of participants in 3D virtual space, we demonstrate a mutual relationship between body movements and fear memories in humans. First, we demonstrate that the acquisition of fear memory results in the adaptation of body movement patterns toward threat-predictive cues. Second, we show that training to actively avoid threats with naturalistic defensive movements leads to a long-term (24 hrs) alleviation of acquired fear memories. This long-term effect was specific to the embodied active-avoidance training, as the less-embodied control training with extinction or vicarious defense only led to a transient reduction of fear which returned 24 hrs later. These results highlight that adapting body movements is a process intrinsic to both acquisition and modification of fear memory among humans. This study thus bridges the previous gap between human fear memories studied in less-embodied experiments versus those experienced in a more-embodied real life.

### Fear acquisition in body movements

Extending the previous animal literature, we show that human participants acquire distinct body movement patterns through threat conditioning. The non-human animal literature postulates body movements as one primary consequence of acquiring fear memories (*11*, *55*). Our study demonstrates direct evidence that humans are not exceptions. In human studies, body movements have been generally marginalized compared with other common measures such as physiological, subjective, or neural responses (*17*, *56*). Although body movements may appear ancient and thus less relevant to modern humans, various threats, such as interpersonal violence and fire, still require one’s whole-body movement for protection. Thus, adapting body movements may serve as a central fear memory function that can directly facilitate survival in humans.

### Fear alleviation through avoidance actions

Our results further show that embodied training to actively defend against a threat has fear alleviation effects, alleviating CRs in physiological responses and body movements in tandem (**Fig 3C**). Crucially, the embodied active-avoidance training was resistant to a spontaneous return of physiological defensive responses (i.e., SCR) with the passage of time, overcoming one of the major constraints of conventional extinction training (*52*, *57*). We thus extend the evidence for the advantage of active-avoidance training, relative to extinction training, in resisting a long-term return of fear (*16*, *25*, *28*–*30*) with our embodied procedures. Furthermore, we demonstrate that the long-term fear reduction effect is improved through the embodiment of active-avoidance relative to the less-embodied vicarious avoidance experience (Group 2). Together, the results advance the potential clinical application of active-avoidance training to overcome traumatic memories for various real-life threats that require naturalistic bodily actions for defense.

### Fear reduction mechanisms with embodied active-avoidance

Unlike the embodied active-avoidance training, fear alleviation effects with our virtual extinction training was transient and followed by the spontaneous recovery of fear responses 24 hrs later. This result is in line with the theoretical and experimental studies suggesting that extinction is a form of inhibitory learning rather than an erasure of acquired fear (*52*, *58*, *59*). The absence of spontaneous recovery with the active-avoidance training suggests that its underlying mechanism may not rely on the inhibitory mechanisms as the conventional extinction training. Supporting this possibility, our preliminary results showed that participants who displayed a larger interim fear reduction effect through the Extinction training resulted in a larger spontaneous recovery 24 hrs later, suggesting that temporary suppression of fear response during the Extinction training may be uninhibited with a passage of time (**Figure S3-2C**). Importantly, the interim fear reduction effect through active-avoidance training showed no such correlation with the spontaneous recovery of the fear response.

The underlying neural mechanisms for the fear alleviation effect through the embodiment of active-avoidance training remains to be studied in future studies. Previous studies with human participants have elucidated the role of the network among the medial prefrontal cortex (mPFC), amygdala, and striatum in the successful performance of active-avoidance (*16*, *26*). Furthermore, studies with rats suggest a more detailed mechanism in which mPFC mediates the trade-off between passive fear-like responses and active-avoidance (*2*, *9*, *60*). Future studies may examine how the embodiment of active-avoidance training augments fear alleviation processes via mPFC and/or other areas in humans. For example, embodied procedures might recruit additional brain areas such as the insular to regulate the fear memory circuit by integrating the bodily feedback signals (*61*).

### Embodied versus vicarious active-avoidance

Our results from Group 2 revealed that participants’ own bodily defense action itself facilitates fear alleviation effects of active-avoidance. Specifically, the embodied active-avoidance training led to a long-term fear reduction only when participants executed the defense action (spraying), while mere observation of a third avatar defending on their behalf did not. Previous studies have demonstrated that both fear acquisition and extinction can occur through the observation of others (*62*–*65*). Unlike the acquisition and extinction of fear memories, our results suggest that fear alleviation through avoidance does require participants’ own actions and vicarious avoidance is not sufficient to reduce fear.

A sense of control or agency over negative events has been suggested as a critical factor on fear responses to stimuli signaling threat (*66*). The fear alleviation effect of the participants’ own defense movement could be due to their body movement, sense of control, or both. As the sense of control is intrinsic to bodily defense movements, the contributions of movement and sense of control may be studied more systematically in the future.

### On clinical application of embodied active-avoidance

Recently, the extinction of fear in VR settings has been applied to clinical settings. A meta-analysis holds that extinction with VR may be slightly more effective than non-VR extinction procedures (*67*). Here, our results suggest that the embodied approach to leverage the VR techniques may potentially improve treatments for anxiety and fear-related disorders such as PTSD. Because a third of patients with PTSD do not respond to conventional exposure therapies (*68*–*70*) which are based on extinction procedure (*71*), the embodied active-avoidance training may be considered as an alternative for those non-responders in the future clinical trials.

Interestingly, we found that male participants were more likely to benefit from the embodied active-avoidance training relative to the extinction training (**Fig 3E**). Preliminary, this suggests that embodied active-avoidance training could be a more effective treatment choice, especially for male than female patients with traumatic memories. However, as the use of only male violent avatars limits our interpretation of the participants’ gender effect, future studies should more directly test the gender effect.

### Limitations

Despite the aforementioned advantage of embodied active-avoidance training over extinction training, some of the limitations should be cautiously considered when potentially applying embodied active-avoidance training for clinical intervention. Specifically, the active-avoidance training transiently led to higher SCR to CS+ than the extinction training when the interim effects were assessed at the end of the training sessions (**Fig 3D**). These results suggest that some individuals might momentarily experience more distress with the active-avoidance training than with the extinction training, even though the active-avoidance training would later result in a better long-term fear alleviation effect. Thus, embodied active-avoidance training may be applied to reduce fear when individuals can readily tolerate those limitations to achieve better long-term fear alleviation effects.

In summary, we demonstrate the critical role of body movements in both acquisition and alleviation of human fear memory. The findings may benefit the field by calling to re-centralize the body in research of human fear and its disorders such as PTSD.

## Materials and Methods

### Participants

Forty-one adults (20 males, Mean age = 25 ± 6 years old) without a reported history of anxiety or related disorders were paid to participate in the three-day experiment (8000 yen per day including agent charge) as Group 1. All participants provided written informed consent prior to their participation. The sample size was estimated based on our pilot study with G*power 3.1.9.6 (*72*). To detect a conditioning effect in our virtual conditioning session (the difference in SCR between CS+ and CS-) with a one-tailed t-Test yielding an effect size of dz = 0.54 with a power (1-β err prob) of 0.95, the required sample size was estimated to be *N* = 39.

The sample size was rounded to 40 participants to ensure counterbalancing. Forty-one participants were enrolled in the end as one participant failed to complete the last session. This participant did not complete the reconditioning session (see Procedures) but the data from other completed sessions were included in the analyses. This sample size is similar to a previous study with active avoidance among 28 human participants (*25*).

Fifty-four adults (27 males, Mean age = 23 ± 3 years old) were additionally enrolled as Group 2. All participants provided written informed consent prior to their participation. Participants in Group 2 were divided into two sub-groups (Self-group: *N* = 27, 14 males, Mean age = 24 ± 2; Vicarious-group: *N* = 27, 13 males, Mean age: 24 ± 3). One participant from the Self-group stopped the experiment due to a malfunction of the equipment and the data was not included.

### Procedures

#### Group 1

The experiment consisted of four main sessions conducted across three consecutive days (**Fig 1A**): Threat-conditioning (day 1; 24 min), extinction (EXT), and embodied active-avoidance (ACT) training sessions (day 2; 15 min each), and spontaneous recovery test (day 3; 11 min). The test session on day 3 was followed by a renewal test session in a novel context and a reconditioning session. All sessions were conducted in virtual settings with a head-mounted VR display (HTC VIVE Pro Eye) (see Materials for details). The study was approved by an IRB committee at Sony (Tokyo).

##### Mirror sessions

Prior to each experimental session, a participant was virtually situated in the back corner of an elevator for one minute to enhance an immersive feeling (*73*). The elevator was identical to the experimental sessions except that a mirror appeared toward the wall on the left side from the participant. A mirror reflected him or herself as a human-like avatar. The avatar-self was semi-personalized to match the gender of each participant. A participant was asked to move around his or her body to observe the corresponding movements of the avatar-self. Only prior to the ACT session, the avatar-self appeared with a spray in the right hand. The position of the hand-held controller was reflected as a virtual defense spray can in the hand of a participant’s avatar-self, and pulling a lever on the hand-held controller resulted in triggering to emit virtual mist from the spray can. The participant practiced for one minute to position their bodies to trigger a virtual defensive spray to exert active-avoidance movements.

##### Experimental sessions

On day 1 participants underwent a threat-conditioning session with two CS+s and one CS- (**Fig 1C**). In the conditioning session, three male avatars served as CSs and appeared on 10 trials each without showing any violence. Two of the avatars appeared on additional 15 trials with violence, hence they both served as CS+s. One avatar never appeared with violence and thus served as a CS-. The assignment of the three avatars to each CS was counterbalanced across participants. The task of participants was simply to observe the avatars as an avatar-self throughout the session. The movements of the participants were reflected in the movement of their avatar self, which was semi-customized to reflect their gender in all sessions. On each trial, a participant was virtually situated in the back corner of an elevator. The trial sequence was as follows (**Fig 1B**, see **Movies S1-3**): After 4 sec, the door of the elevator opened and one out of the three male avatars CSs entered the elevator after which the elevator door closed. In safe trials without violence, the avatar idled with his body facing toward the participant for 2 s and then turned around to face the closed door to remain standing semi-still for the rest of the trial (8 sec). On the trials with violence (CS+), an avatar entered the elevator and idled with identical movements as in safe trials (CS-). However, after 6 s the avatar quickly walked toward the participant and raised his right arm to hit the participant’s head. Each trial lasted 15 sec and inter-trial interval was 6 sec.

The experience of being hit in the realistic VR environment served as threatening unconditioned stimuli (US). The hitting action by the aviator was accompanied by a corresponding sound effect. The opening of the elevator door was also accompanied by a sound effect and the background machinery noise of the elevator was constantly played throughout each trial to enhance a realistic VR experience. Three avatars were all males to reflect the relatively higher prevalence of violent incidence by males and related psychological traumas (*36*).

The trial order was pseudorandomized with the restriction that no more than two consecutive trials of the same condition occur (e.g., a male avatar A with violence) to avoid habituation (*17*). Each of the initial two trials was always with one of the two CS+s avatars with violence in a counterbalanced order across participants. This was to boost expectation for threat in the session (*74*) and to avoid inducing any difference in potential *blocking* of conditioning effects (*75*, *76*) due to unequal numbers of safe trials with the two CS+s prior to the occurrence of initial violent trials.

On day 2, participants went through two training sessions: the embodied active-avoidance (ACT) and Extinction (EXT) sessions. The order of the two sessions was counterbalanced across participants and was separated by a 15 min break. Only one of the CS+s appeared during the ACT session (CS+_active_) and only the other CS+ appeared during the EXT session (CS+_extinction_), along with the CS- in both sessions.

In the ACT session, CS+_active_ and CS- appeared 25 and 10 times, respectively. Among the trials with CS+_active_, CS+_active_ appeared 15 times showing the common entering and idling actions followed by preparatory actions for hitting (i.e., approaching a participant to raise an arm over the participant) as in the first 9 sec of the CS+s trials in the conditioning session (**Fig 1**). However, here CS+_active_ trials never ended with actual hitting due to the preventative defense of spraying the CS+_active_ avatar by participants. In the remaining 10 trials, CS+_active_ showed no sign of violence as in the trials with CS-. Those trials without violence were included to equate the CS-US contingency rate between the conditioning session on day 1 and the ACT sessions on day 2. Prior to the ACT session, participants were instructed to use a controller in their right hand as a virtual defense spray against the CS+_active_ avatar upon seeing his preparatory actions for hitting (i.e., approaching and raising a hand). Participants positioned their bodies to target the avatar then pulled a trigger on the controller to spray the white mist toward the CS+_active_ avatar. Upon the spraying, the avatar displayed suffering in response to the released spray with his hands covering his face (**Fig 1B**, see **Movies S2**). The spraying action was practiced for one minute during the mirror session prior to the ACT session (see *Mirror session*). Thus, only the CS+ but not the US (hitting) appeared during the active control session.

In the EXT session, the CS+_extinction_ avatar appeared 25 times with the same action as the CS- avatar (i.e., no violence) (CS+_extinction_) along with CS- appearing 10 times. Thus, the total number of trials (35) was identical to that of the ACT session. In the EXT session, participants were simply instructed to observe the avatars.

The order of the trials in the ACT and EXT sessions was pseudo-randomized while avoiding three consecutive trials with the same condition. The context of the two training sessions was the identical elevator space as in the conditioning session. In the ACT session, the first two trials were always without precursor actions for violence to avoid the overestimation of the violence frequency in the upcoming trials (*74*) relative to the EXT session.

Participants underwent the spontaneous recovery test on day 3. CS+_active_, CS+_extinction_ and CS- avatars appeared 8 times each. The trial order was pseudorandomized with the first trial always being CS- to capture an irrelevant orienting effect (*25*, *77*) and its data was not included in the analysis. The participants did not hold the virtual security spray during the sessions on day 3.

Participants then underwent an additional test session with a new context (dimly lit virtual elevator) and reconditioning session. As the conditioned responses were diminished through extinction during the unreinforced trials from the spontaneous recovery test, the details of those extra sessions and their results are presented in Supplementary (**Fig S5**).

##### Rating sessions

Participants provided subjective ratings on their fearful feelings and US (violence) anticipation upon seeing the CS avatars in the back corner of a virtual elevator. Throughout the experiment, they provided ratings 6 times in total; before and after the conditioning session, after the active-avoidance, after the Extinction sessions, after the renewal test, and after the reconditioning session. In each rating session, the three CS avatars appeared once in a counterbalanced order across participants. Each avatar entered the elevator and paused with the panel appearing in front of him to display a question “how fearful do you feel upon seeing the man?”, followed by another question “how likely do you think the man is going to hit you?” in this order. Participants provided verbal reports of their ratings on a seven-point scale, and an experimenter recorded the ratings.

#### Group 2

We conducted the experiment in Group 2 across two consecutive days. The conditioning session was identical to Group 1, and some modifications were introduced only during the ACT training sessions (day 1) and spontaneous recovery tests (day 2).

To examine whether the alleviation of conditioned fear depends on the embodiment of the ACT training the second group of participants with body motion tracking (**Fig 1F**) went through the ACT training with (Self-group) or without their own movements (Vicarious-group). While overall procedures were similar to Group 1, there were some exceptions. First, since the aim was to compare the alleviation of fear responses to CS+_active_ between the groups, we did not include the EXT training as a reference. Instead, participants underwent the ACT training with one of the two avatars (CS+_active_) while the other CS+ received no training (CS+_untrained_). Second, both conditioning and ACT training sessions were conducted on day 1.

##### Embodied active-avoidance defense training sessions

In the ACT training session, participants in the Self-group underwent the same procedure as in Group 1, where they actively defended themselves with a virtual security spray to prevent violence. Participants in the Vicarious-group observed a third-passenger avatar defending against the violent avatar with a security spray on their behalf. In both groups, the third avatar accompanied the participants in the training sessions to equate the potential effect of social presence in fear responses (*78*) with the Vicarious-group. The third avatar was a female passenger that stood in front of the participant’s standing position, facing the center of the elevator, in front of the number button on the right side of the elevator door (**Fig 4B**). The gender of the third avatar was fixed to be female across participants to make her clearly distinguishable from the CS male avatars so that participants would not potentially anticipate threat (violence) from the third avatar. When the male avatar CS was going to attack in the Vicarious ACT training session, the third avatar sprayed the virtual security spray toward the head of the avatar at the time when the paired Self-group participant sprayed on the same trial. When the avatar CS did not inflict violence, the third avatar operated her smartphone to reflect a common encountering scene with others in an elevator. The third avatar, however, lowered her smartphone and looked up at the male CS avatar when the participants in the Self-group initiated the spraying action, as the absence of reaction from the third avatar would otherwise appear unrealistic. Since subjects did not undergo EXT training, one CS+ was associated with ACT training (CS+_active_) and the other CS+ was untrained (CS+_untrained_).

##### Spontaneous recovery tests

Participants underwent the spontaneous recovery test as in Group 1. They additionally underwent a second test and reconditioning sessions. As the conditioned responses were weakened through extinction during unreinforced trials from the first spontaneous recovery test, the details of those sessions are presented in Supplementary (**Fig S6**). Briefly, we conducted the second test in the same context as in the conditioning session but with a third avatar (**Fig 4B**) from ACT training sessions to examine the effect of social presence on fear memory retrieval. Participants were informed that a passenger would appear prior to the second test.

##### Rating sessions

Participants in Group 2 provided subjective ratings in the similar manner as Group 1 for 6 times in total; before and after the conditioning session, after the active-avoidance defense sessions, after both the first and second spontaneous recovery tests and after the reconditioning session.

### Materials and equipments

The interactive VR images were programmed with the Unity 2019.3.0 and were presented at 60 Hz with a head mounted display, the HTC VIVE Pro Eye (2019, https://www.vive.com/eu/product/vive-pro-eye/overview/). The tracking of head positions (x, y, z) were also enabled by the HTC VIVE Pro Eye and the tracking of hand positions (x,y,z) were obtained by the Vive Trackers bound to the right hand. The image presentation and the recording of the tracked data were controlled by a laptop (G-Gear: N1588J-710/T) with NVIDIA^®^ GeForce RTX^TM^ 2070, Intel^®^ Core^TM^ i7-9750H processor, and Windows 10 Pro. Eye-tracking was calibrated with the Super Reality (SR) runtime implemented in the HTC VIVE Pro Eye prior to each experimental session. Their skin conductance was measured by Shimmer3 GSR (LoAndStream v0.11.0, Shimmer^®^) with its electrodes attached to the participants’ left palm. The skin conductance signal was recorded at 250 Hz with a Tobii Pro Lab software (ver 1.145.28180) installed in another laptop (Dell: Precision 7720) with Windows 10 Pro.

### Body Movement

Positional and rotational data was obtained by the Vive Trackers and both head and hand motions of the subject in the VR were continuously tracked at 60 Hz during all experimental sessions in both Group 1 and 2. We obtained head and hand positions and rotations across the three axes (x, y and z) (**Fig 1E)**. The sensor measurements of the trackers are provided in a head-fixed coordinate system where the x-axis points to the right, the y-axis points upward and the z-axis points forward.

### Analysis

#### Subjective rating

Subjective ratings on fearful feelings and US anticipation were analyzed separately. Differential subjective ratings for both CS+s were calculated relative to the CS- within the same rating sessions (i.e., subjective CRs). Pre-conditioning ratings were used as baseline to correct ratings in each session. Exceptionally for the two post-training rating sessions, subjective ratings were analyzed for CS- and only one of the CS+s which appeared during the training session immediately before the rating session. This was to avoid including the rating data for the untrained CS+ due to the counterbalanced order of the two training sessions across participants.

#### SCR

SCR data were low and high-pass filtered (0.015 - 0.05 Hz) with an FIR filter, detrended and normalized. The SCR onsets and peaks were automatically detected by Tobii Pro Lab software (Tobii Pro AB (2014). Tobii Pro Lab [Computer software]. Danderyd, Sweden: Tobii Pro AB.). The cumulative SCR was computed as the summation of all the through to peak amplitude deflections of individual SCR(s) within each trial, to effectively quantify the responses during the relatively long trial (14s) with a naturalistic VR scene with dynamically changing CS avatars. Only when the peaks and corresponding onsets of individual SCR(s) were observed within a trial, their values were accumulated as they may otherwise reflect irrelevant responses such as anticipation for the start or end of a trial. The cumulative SCR was log-transformed and scaled to the maximum cumulative SCR of the CS trials (not followed by US) of the conditioning session. For the conditioning and reconditioning sessions, the CS trials with US were not included in the analysis to exclusively examine the conditioned responses. The test sessions always started with one CS- presentation to capture the orienting response and was not included in analysis.

### Movement Representational Similarity Analysis

#### Tracking of 2 body parts with VR system

To investigate the relationship between the patterns of participants’ body movement and learned threat associations we used Representational similarity analyses (RSA) (*45*–*47*). We calculated the similarity of body movements upon seeing different CS avatars based on the position and rotation of both head and hand for the three axes (12 dimensions, see **Fig 1D**). Note that the trials with US (violence) were removed from the analyses so that the movement pattern differences, if any, would reflect the movement deviations in anticipation of potential violence. To obtain a measure that is independent of coordinate values and unbiased by previous positions during a trial, we calculated the absolute difference between two consecutive recorded samples (|Position(t2)-Position(t1)|) within trials. This yielded a measure of sample-by-sample change in the movement for each of the 12 movement parameters. For all analyses, CS+s trials with a US (violence) were discarded, such that the remaining CS+s trials were identical to the CS- trials except for the avatar identity (CS+_active_, CS+_extinction_, or CS-, see **Fig 1C**). Next, to reduce the number of comparisons we divided the data within a trial into 10 time-windows of 833 ms (50 samples) each. We created a vector per condition (e.g., CS+ or CS-) for each participant per time window. Each vector now contained the spatial pattern of movement (12 movement parameters) within a specific time window (50 time-samples), related to a particular condition. Pairwise Pearson correlations were calculated between vectors of condition pairs, resulting in a similarity measure, depicting correlations among conditions for each participant per time window. For instance, the correlation between movement in response to the CS+s and CS- avatar was provided for each time window of the trial, and for each participant a metric of how similar the movement in (non-violent) CS+s trials was relative to CS- trials. Correlation coefficients were then Fisher-transformed and compared using classical and bayesian paired t-tests.

The RSA was first conducted to assess threat conditioning by comparing the CS+s to CS- similarity between the early and the late phases of the conditioning session (trials 1-2 versus trials 9-10, one-sided paired t-tests). We expected the CS+s to CS- similarity to decrease in the late phase of the conditioning session due to the emergence of threat conditioning effects in body movements. Second, RSA was performed to compare the effect of embodied Active-avoidance and Extinction training on spontaneous recovery on day 3 by comparing CS+_active_/CS- similarity versus CS+_extinction_/CS- similarity in the first two trials of the test session. We expected the ACT training to be more effective in reducing the fear response, resulting in greater CS+_active_/CS- similarity compared to CS+_extinction_/CS- similarity. For the RSA analyses, we used the NeuroRA (https://neurora.github.io/NeuroRA) toolbox (*46*).

#### Tracking of 31 ROIs with motion tracking system

Positional and rotational data during Experiment in Group 2 were additionally obtained by Motive OptiTrack Flex13. A set of 39 markers were distributedly placed based on the conventional body coordinate of OptiTrack covering each participant’ body surface (**Fig 1F**). The position on the 3D axes of 39 markers were tracked. Four markers for a head were placed on the nearest surface of the VIVE headset to keep the markers visible for tracking. Eight OptiTrack Flex13 cameras captured the motion of 39 markers and were recorded with the OptiTrack Motive 2.3 software at 120 Hz. The recorded motion data were preprocessed with OptiTrack Motive 2.3 to fix body-position labelings of markers when their labels were swapped between markers during recording or when their labelings were lost for some data points. Gaps in motion tracking data corresponding to the time points where marker(s) were disattached or hidden by participants’ body parts from the tracking cameras during less than 10 s were linearly interpolated with Matlab. Six out of 39 markers were discarded from the analysis because data was lost in 30 - 42% of the participants, and we conducted subsequent analyses with the remaining 33 markers. Additional markers with gaps longer than 10 s were discarded from one session in five participants.

The RSA analysis in Group 2 was based on 33 markers covering both sides of the body (**Fig 1F**). For each body marker, we systematically grouped 2 nearest markers (including both sides of the body) to form a ROI clustering 3 markers, to examine the local, yet spatially distributed, patterns of movement. Avoiding overlaps of clusters with identical 3 markers, this procedure yielded 4 ROIs in the head, 8 ROIs in the trunk, 11 ROIs in the arms and 8 ROIs in the legs (31 clusters in total).

### Statistical Analysis

Where Shapiro-Wilk’s test indicated that normality assumptions were violated, Wilcoxon correction was applied. In addition to traditional null hypothesis significance testing (NHST), we computed Bayes factors (BF) for all analyses. BF allowed us not only to find evidence for our tested hypotheses but to quantify the evidence in favor of the null hypothesis. We based our results on BF especially when it was beneficial to infer the relative degree of evidence across multiple time windows (10) within a trial to capture the temporal dynamics in the body movement analyses. For the Bayesian analyses, a Cauchy prior distribution centered around zero was used with an interquartile range of r = 0.707. All analyses were performed using JASP (Version 0.16).

We used paired one-tailed t-tests and directional Bayesian t-tests (BF_+0_) in examining the predicted conditioned effect (i.e., larger responses to CS+s than to CS-) and the more robust long-term (24 hrs) fear reduction effect of the ACT training than the EXT or no-training, as predicted based on previous literature (*25*). Otherwise, we used two-tailed t-tests and non-directional Bayesian t-tests (BF_10_). To test for evidence supporting the null hypothesis, we performed Bayesian paired samples t-tests (BF_01_). We used a common interpretation of the Bayes factors (BF 1-3: anecdotal evidence, BF 3-10: moderate evidence, BF>10: strong evidence) (*79*). For the classical t-tests of body movement similarity conducted across 10 time windows, we corrected for multiple comparisons by fixing the false discovery rate (FDR) at 0.05 (*80*).

We did not exclude any participants based on the absence of differential SCR to CS+s relative to CS- during the conditioning because those with less differential CR could be more prone to anxiety and thus represent a relevant cohort in fear memory research (*20*, *44*). In addition, we did not use SCR as exclusion criteria given that we aimed to evaluate the degree of threat-conditioning multidimensionally, not only with SCR but with additional measures of bodily movements and subjective ratings. However, we shared how results remain quantitatively similar after removing SCR-based outliers across sessions. The changes in SCR and movement data within or across sessions were analyzed by examining responses in the early and late phases (two trials each) within a session. As the subjective ratings were collected only once after each session type, the data were not divided into smaller phases.

To examine the effects of participants’ gender and training order on fear attenuation in Group 1 and the effects of ACT training type (Self-versus Vicarious-group) on fear attenuation in Group 2 we used Repeated Measures ANOVA and Tukey test for multiple comparisons.

## Acknowledgments

We thank Yuna Ujiie and Kazuto Ishii for their support in conducting experiments for their support with the equipment. We appreciate Anikó Kusztor, Ben Seymour, and Jessie Taylor for their helpful comments on the manuscript.

## Funding

Japan Science and Technology Agency (JST) Presto (18068712) (AK)

Japan Science and Technology Agency (JST) Moonshot (20343198) (AK, KN, KW)

Japan Society for the Promotion of Science (JSPS), Grant-in-Aid for Scientific Research (B) (18H02714 and 22H01111) (AK).

## Author contributions

Conceptualization: MAG, AK, TC, HI

Methodology: MAG, MEW, TN, KW, KN, HY, NK

Investigation: MAG, MEW, AK

Visualization: MAG, MEW

Supervision: AK, HI, KN, KW

Writing—original draft: MAG, AK, MW

Writing—review & editing: HI, TC

## Data and materials availability

All data needed to evaluate the conclusions in the paper are present in the paper and/or the Supplementary Materials.

## Supplementary

### Supplementary Text

#### Extension of Fig. 1B

Supplementary videos depicting scenes from the trials with CS+ (US), CS+ during ACT training, and CS+ (no US) during EXT training are captured in **Movies S1-3**. The windows 1-10 as well as epochs (eg, US anticipation) are labeled within the movies. The sounds are not included for quality.

**Movie S1**: CS+ (US)

**Movie S2**: ACT training

**Movie S3**: CS+ (no US) or EXT training

#### Additional statistical results Fig. 2D

In addition to the ROIs in the right arm and feet described in the main text, we merely observed anecdotal evidence of CRs for ROIs in the head and trunk in the *reveal-* and *violence-epoch* [Head: time window #1: 1 ROI BF_+0_=1.61; Trunk: time window #1,10: 3 ROIs BF_+0_>1.16; all one-tailed]. Here, one-tailed tests were used to restrict the results with positive conditioned responses (i.e., decrease in similarity between CS+ and CS-) for interpretation. The bodily topographical maps additionally depicted several ROIs with increased movement similarity between the CS+_active_ and CS- encounters from the early to late phase of the conditioning session. The increase of similarity could reflect the generic reduction of movement variability due to the learning of the spatio-temporal structure in the scenes, confining the participants’ bodily movements such as where they would attend to in the 3D space in predicting the next move of the avatar. To capture these increased similarity in movements, we also ran one-tailed t-tests. Bayesian factors indicated that movement similarity in legs during CS+_active_ and CS- encounters increased from the early to late phase of the conditioning session across several windows [time windows #1-5, 7 & 8: 2 ROIs moderate evidence BF_+0_ >5.62; 2 ROIs anecdotal evidence BF_+0_ >1.03].

#### Extension of Fig. 3: Training effects on subjective ratings

Interim and long-term fear reduction effects were also assessed with subjective ratings (**Fig S4**). We note that capturing spontaneous recovery of fear, if any, would have been difficult with subjective ratings. This is because, despite the fact that spontaneous recovery typically emerges only on initial few trials after time lapse and swiftly disappears (50), participants provided their ratings only after completing the two long-term test sessions (spontaneous recovery test and renewal test) each composed of 24 trials. The experimental design to initiate ratings only after experimental sessions was to avoid interfering with the measurement of spontaneous recovery in SCR.

#### Statistical results for Fig. 4C

Bodily topographical maps depicting ACT training effects relative to no-training captured distinctive spatio-temporal patterns of the training effects between the Self- and Vicarious-groups in Group 2. In addition to the training effects captured in the arms of the Self-group during the *Violence-epoch* by moderate and strong evidence described in the main text, Bayesian statistics also revealed effects in the arms and trunk in the same time windows with anecdotal evidence [Arm’s ROIs: time window #8: 2 ROIs BF_+0_>1.25; Trunk’s ROIs: time window #8: 2 ROIs BF_+0_>1.14; all one-tailed]. Self-group additionally showed a training effect in the right arm during *Idling-epoch* as indicated by anecdotal evidence [time window #4: 5 ROIs BF_+0_>1.10]. Additionally, Self-group showed training effects in non-threat relevant windows (*reveal-epoch*) in left arm and leg as indicated by anecdotal evidence [Left arm: time window #1: 3 ROIs BF_+0_>1.24; Left leg: time window #2: 1 ROI BF_+0_=1.86; all one-tailed]. The Vicarious-group showed anecdotal effects in spatially disconnected ROIs within the *reveal-, idling-* and *anticipation-epochs* but no effect in *violence-epoch* [Arm’s ROIs: time window #1: 3 ROIs anecdotal evidence BF_+0_>1.18; Trunk’s ROIs: time window #3,4 & 5: 3 ROIs BF_+0_=1.05; Leg’s ROIs: time window #3,6 & 7: 2 ROIs BF_+0_>1.25; Head’s ROIs: time window #4 & 6: 3 ROIs BF_+0_>1.03; all one-tailed].

#### Statistical results for Fig. 4D

SCR toward both CS+ avatars was statistically similar between the two groups before the ACT training session. Repeated measures ANOVA with one between-subjects factor of training type (Self and Vicarious training) and one within-subjects factor of CS+ type (CS+_active_ and CS+_untrained_) showed that there was neither a significant main effect of ACT training nor an interaction between ACT training type and CS+ type in the conditioning session [ACT training type: F(1, 50) = 0.77, *p* = 0.38; ACT training type * CS+ type: F(1, 50) = 1.56, *p* = 0.21]. Repeated measures ANOVA on CR towards CS+_active_ (CS+_active_ - CS-) in the early ACT training session also showed no effect of ACT training nor interaction [ACT training type: F(1, 50) = 0,50, *p* = 0.48; ACT training type*CS type: F(1, 50) = 3.48, *p* = 0.06]. The main effect of CS+ was also non-significant [conditioning session: F(1, 50) = 0.04, *p* = 0.82; early ACT training session: F(1, 50) = 2.91, *p* = 0.09], indicating a statistically similar level of CR to both CS+_active_ and CS+_untrained_ prior to the training sessions. The CR to both CS+s [CS+ active: M = 0.020 ± s.e. 0.017; CS+_untrained_: M = 0.007 ± s.e. 0.014], however, were not significantly different from zero (i.e., not significantly larger than CS-) during the conditioning session on day 1 [CS+ _active_: *t*(51) = 1.16, *p* = 0.12; BF_+0_ = 0.49; CS+_untrained_: *t*(51) = 0.48, *p* = 0.31; BF_+0_ = 0.23). During the early ACT training session on the same day 1, CR to CS+_active_ [M = 0.067 ± s.e. 0.097] was also no significantly different from zero [*t*(51) = 0.69, *p* = 0.24; BF_+0_ = 0.28]. CR emerged only from day 2 (**Fig 4D**) potentially due to overnight enhancement of fear memory through consolidation (40-42). As in Group 1 we were inclusive with all participants to reflect critical population diversity in threat conditioning effects (20,44).

---

**Figure S1.**
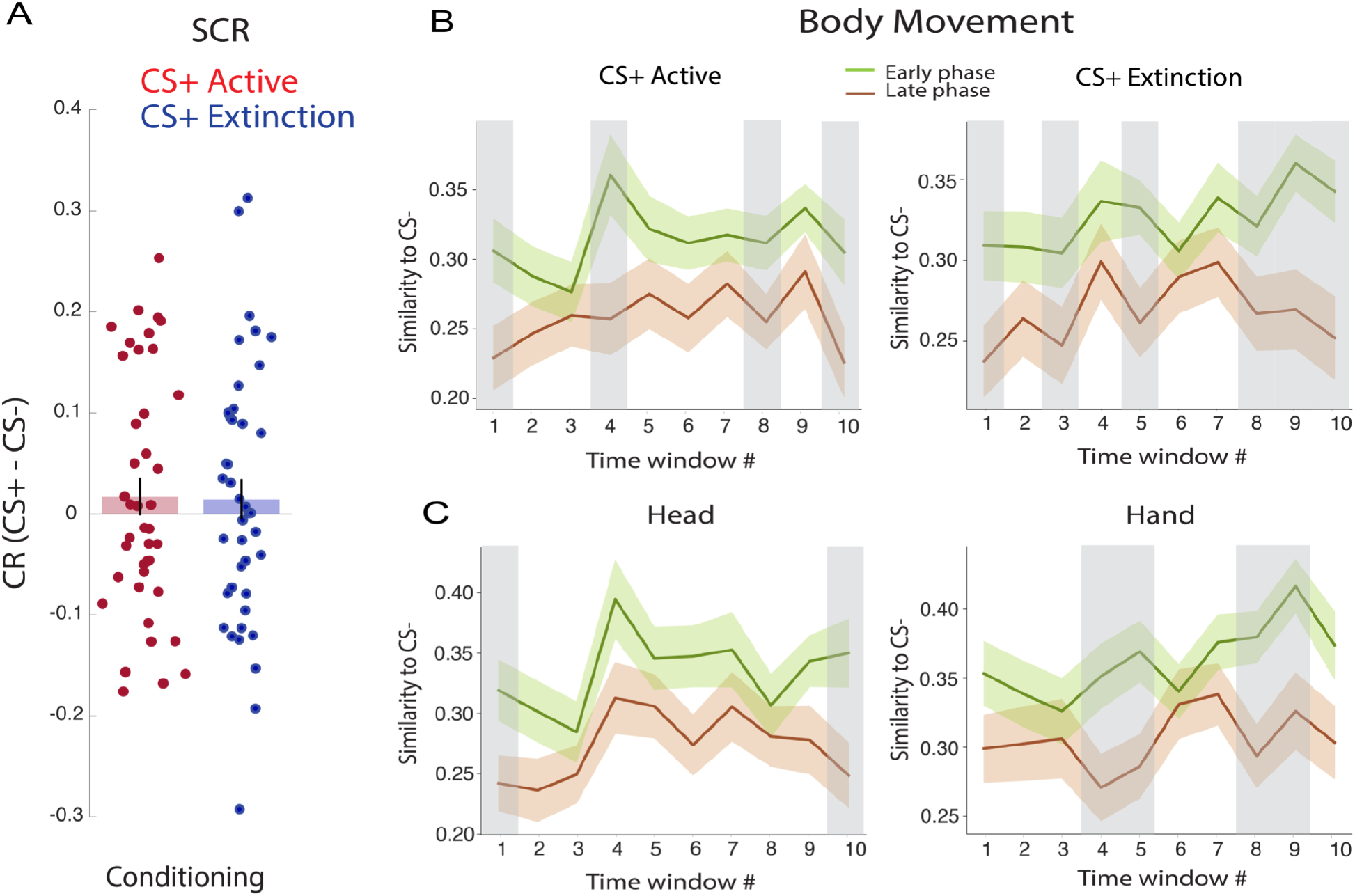
(**A**) Conditioned response (CR) in SCR measures during the conditioning session on day 1. (**B**) The decreasing similarity in body movement patterns between each CS+ avatar (CS+_active_ or CS+_extinction_) versus the CS- avatar encounters. (**C**) The decreasing similarity in head and hand movement patterns between both CS+ avatars versus the CS- avatar encounters from early to late phase of the threat conditioning session. The shaded areas indicate moderate and strong evidence (BF > 3). Data are represented as mean ± between subjects SEM.

**Figure S2.**
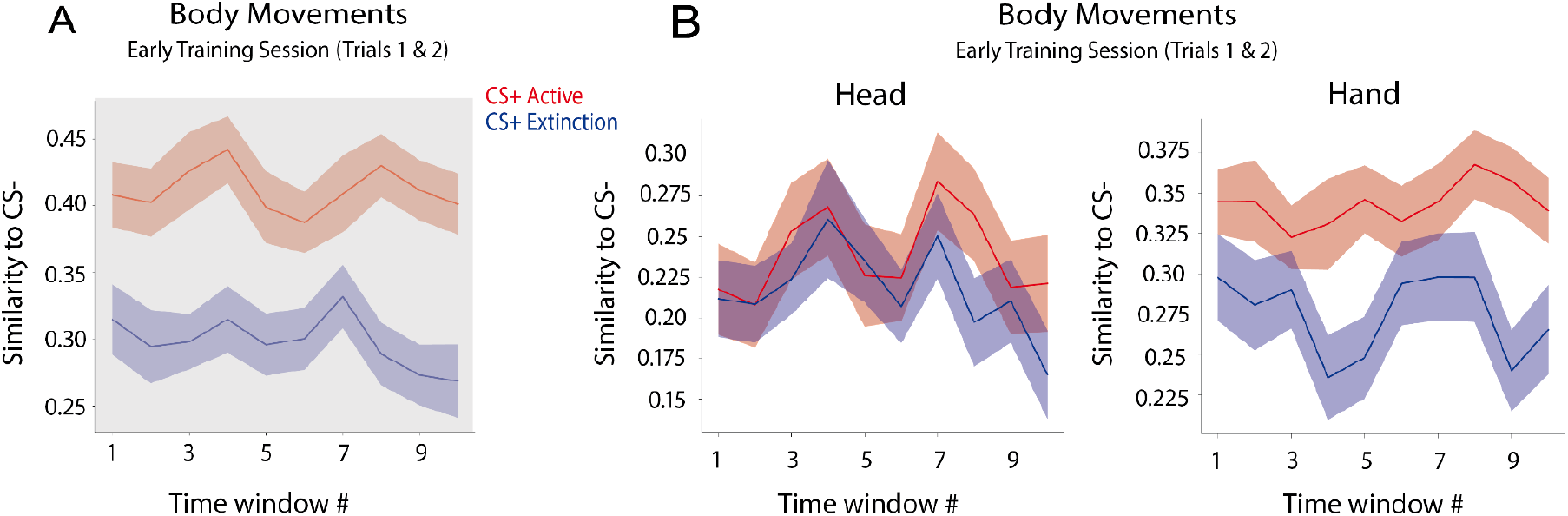
Body movement patterns during the ACT training session showing distinct effects due to the task instruction for action preparations only during the ACT training. **(A)** Already in the early phase of the training (trials 1-2), similarity in body movement patterns during the CS+s versus the CS- avatars encounters deviated between the ACT and EXT condition in all time windows (all two-tailed t-values(40)> 6.43, p-values < 0.001; all BF_+0_ > 25); **(B)** The movement pattern similarity for hand and head separately corroborated that different task requirements (i.e., using a hand-held spray in the ACT condition) confounded the movement pattern analysis especially of the hand during training. Because the task instruction led to apparent differences in body movement patterns already from the start of the training sessions, we did not analyse the interim fear reduction effects during the training sessions.

**Figure S3.**
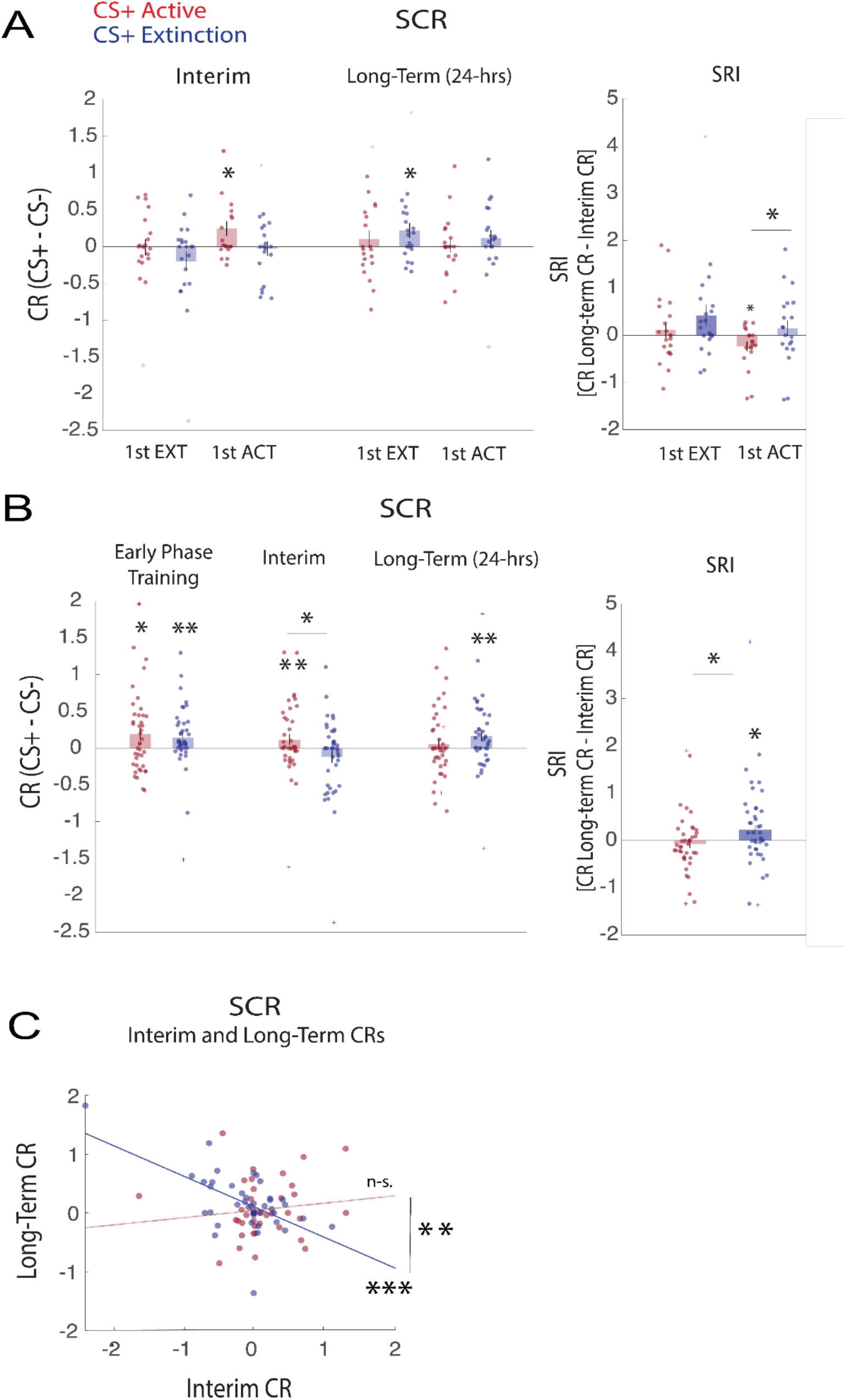
**(A)** Effects of training order on ACT and EXT training. Interim and long-term effects (left) and on SRI (right) of ACT and EXT training separately for Group 1 participants who underwent the EXT training first (EXT-1st: N = 21) and for those who underwent the ACT training first (ACT- 1st: N = 20). We found no interaction between the training order and the CS+ type (CS+_active_/CS+_extinction_) `in both interim [F(1,39) = 0.17、` *p* `= 0.67] and long-term effects [F(1,39) = 0.006、` *p* = 0.93]. *\* correspond to paired t-tests.* **(B)** Conditioned responses in SCR after removing outliers (>=3SD, n = 3, indicated with + signs) from Group 1. Conditioned responses in SCR during the early phase of the training sessions and interim and long-term effects of ACT and EXT training (left) [early phase of the training session: CS+ > CS-; CS+_active_ *t*(39) = 1.97, *p* = 0.02; BF_+0_ = 1.84; CS+_extinction_ *t*(39) = 3.22, *p* = 0.01; BF_+0_ = 26.42; all one-tailed; CS+_active_ Vs CS+_extinction_ *t*(39) = 0.39, *p* = 0.69; BF_01_ = 5.44; interim effects: CS+ > CS-; CS+_active_ *t*(39) = 2.48, *p* = 0.009; BF_+0_ = 5.06; CS+_extinction_ *t*(39) = -0.907, *p* = 0.81; BF_+0_ = 0.09; all one-tailed; CS+_active_ Vs CS+_extinction_ *t*(39) = 2.23, *p* = 0.03; BF_01_ = 3.09; long-term effects: CS+ > CS-; CS+_active_ *t*(38) = 0.89, *p* = 0.18; BF_+0_ = 0.40; CS+_extinction_ *t*(38) = 3.03, *p* = 0.002; BF_+0_ = 16.72; CS+_active_ Vs CS+_extinction_ *t*(38) = -1.26, *p* = 0.10; BF_01_ = 0.08; all one-tailed]. Larger spontaneous recovery index (SRI) after the EXT than the ACT training, as measured by the difference between the interim versus the long-term conditioned SCR [SRI CS+_active_ t(38) = 0.903, p = 0.81; BF_+0_ = 0.09; SRI CS+_active_ t(38) = -2.19, p = 0.017; BF_+0_ = 2.84; all one-tailed; SRI to CS+_extinction_ versus CS+_active_; t(38) = -2.32, p = 0.01; BF_+0_ = 3.70]. All outliers were defined based on the CR values [CS+ versus CS-]. **(C)** Correlation between suppression of CRs through ACT and EXT training (interim) and return of CRs 24 hrs later (long-term). Scatter-plots with least-square regression lines of the intra-subject correlations between CRs across days. Data are represented as mean ± between subjects SEM. *\*p < 0.05 **p < 0.01 ***p < 0.001*

**Figure S4.**
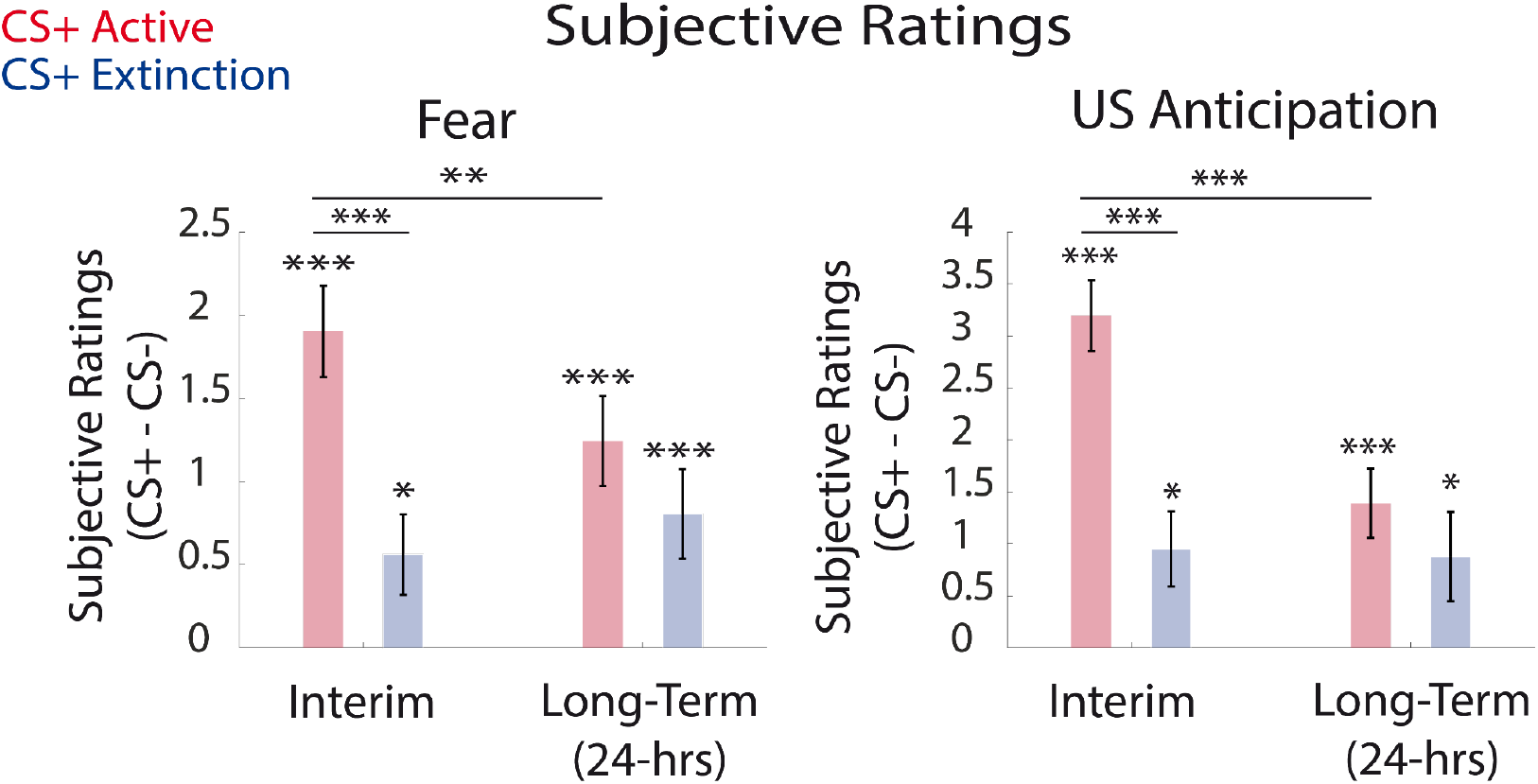
Comparison of subjective ratings effects between ACT and EXT training. Interim (day 2) and long-term (day 3) effects of ACT and EXT training on conditioned responses [CRs: CS+ versus CS-] in subjective ratings of fear and US-anticipation. The post-training subjective ratings to the CS+_active_ were significantly higher than the CS+_extinction_ on day 2 [Fear: t(40) = 4.74, p < 0.001; BF_10_ > 100; US-likelihood: t(40) = 7.38, p < 0.001; BF_10_ > 100]. These results are in line with the SCR results (Fig 3D left) which showed the advantage of EXT training in interim fear reduction effects. This interim advantage of EXT training may reflect anticipation for the precursive actions for violence (US) by the CS+_active_ avatar during ACT training. On day 3, the post-test subjective ratings of fear to CS+_active_ and CS+_extinction_ were no longer different [t(40) = 1.72, p = 0.09; BF_10_ = 0.65]. Fear ratings to the CS+_active_ significantly decreased from day 2 (post-training) to day 3 (post-test) [t(40) = 3.302, p = 0.002; BF_10_ = 16.2] while the fear ratings to the CS+_extinction_ on day 3 remained equivalent to day 2 [t(40) = -0.96, p = 0.34; BF_10_ = 0.26]. This is also in line with the SCR results (Fig 3B) indicating the emergence of the ACT training effect of reducing fear 24 hrs after the training. Accordingly, post-test subjective ratings of US-likelihood to CS+_active_ and CS+_extinction_ were no longer different on day 3 [t(40) = 1.36, p = 0.17; BF_10_ = 0.39]. Rating of US-likelihood for CS+_active_ was decreased on day 3 relative to day 2 [t(40) = 6.89, p < 0.001; BF_10_ > 100], supporting the long-term effects of ACT training. There was no such reduction with CS+_extinction_ from day 2 to day 3 [t(40) = 0.23, p = 0.81; BF_10_ = 0.17]. Together, these results suggest that ACT and EXT training had a comparable long-term effect on subjective fear and anticipation of violence on day 3. Note that CRs (CS+ versus CS-) after both ACT and EXT training were significant on day 2 and 3. Values are baseline-corrected with pre-conditioning ratings as in Fig 2A. Data are represented as mean ± between subjects SEM. \**p* < 0.05 \*\**p* < 0.01 \*\*\**p* < 0.001.

**Figure S5.**
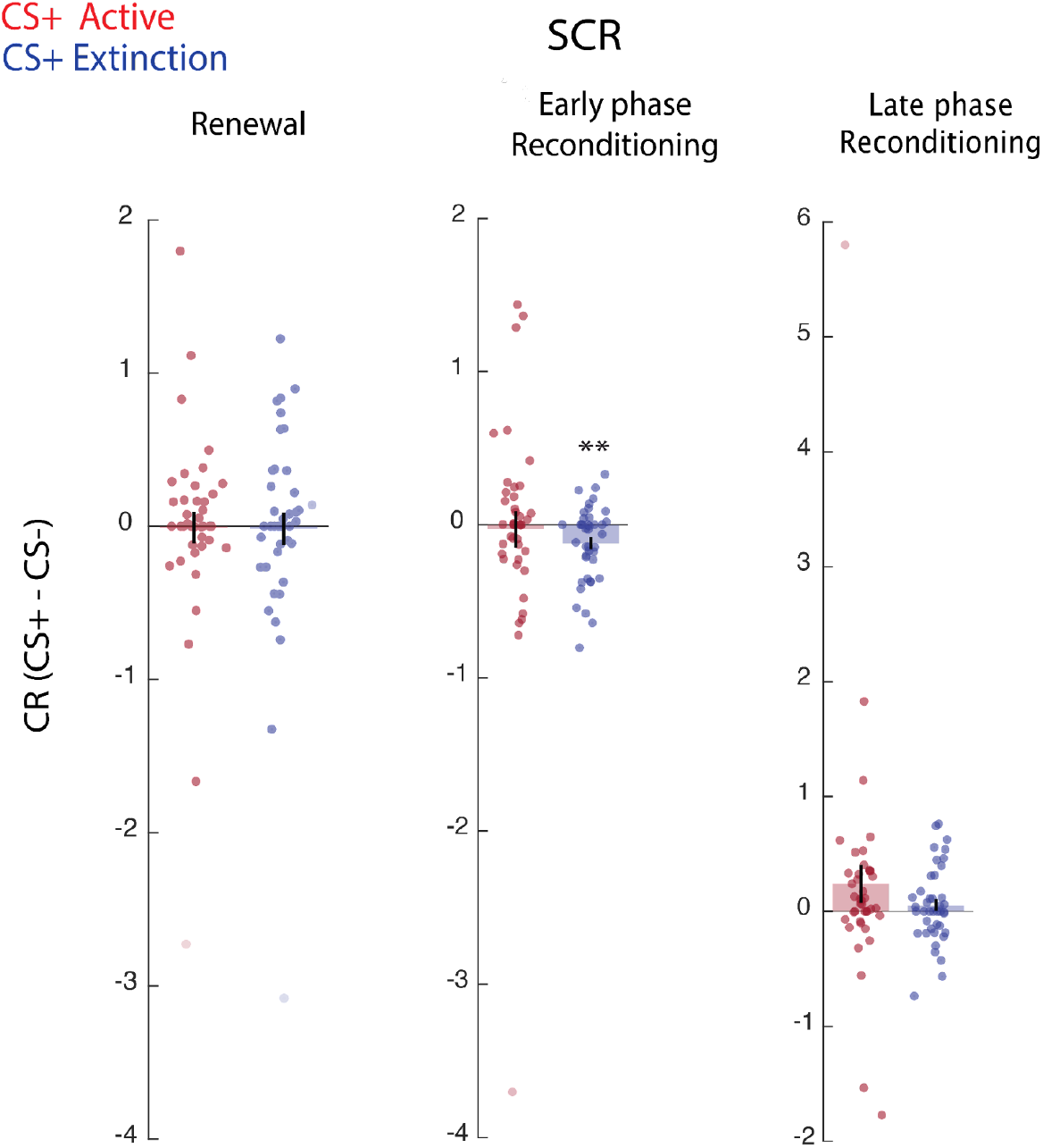
SCR results for the Renewal Test and Reconditioning sessions in Group 1. Day 3 consisted of two test sessions and one reconditioning session. The first test, referred to as the spontaneous recovery test, was performed in the original context as the conditioning session on day 1 and the training sessions on day 2. Next, to examine the effect of context on fear memories (81), a Renewal test was performed in a novel context. The novel context consisted of the same elevator space but with the light drastically dimmed. In both test sessions, the three CS avatars appeared eight times each without violence (US). Finally, participants underwent a reconditioning session to investigate whether the training sessions were effective in reducing the susceptibility to repeated conditioning (82, 83). In the reconditioning session, the CS- avatar appeared eight times without violence, whereas the two CS+s avatars (CS+_active_, CS+_extinction_) appeared eight times without and eight times with violence. The context of the reconditioning session matched the original context. The first two trials of the reconditioning session were always with the two CS+s with violence in a counterbalanced order across participants to resemble the original conditioning session. After the spontaneous recovery test, we also examined whether ACT and EXT training had fear reduction effect when participants were reexposed to the CS+s in a renewal test with a novel context (drastic dimming of lights in the virtual elevator) as well as when they again underwent CS-US pairings in the reconditioning session both on day 3. Participants did not show any CR to the CS+_extinction_ and the CS+_active_ in both the early or late phases of the Renewal Test, and CR did not differ between the two CS+s [CS+_extinction_ Vs CS+_active_; early phase, *t*(40) = 0.15, *p* = 0.87; BF_+0_ = 0.17; late phase, *t*(40) = 0.74, *p* = 0.46 BF_+0_ = 0.21] (Note that only CRs in the early phase of the Renewal Test is shown in the left panel, because later trials are subject to extinction effects due to repetitions of unreinforced trials). This suggests that unreinforced trials in the spontaneous recovery test in the original context extinguished CR to both CS+s before the renewal test. In the reconditioning session, participants showed statistically similar levels of CR to the CS+_extinction_ and CS+_active_ [*t*(40) = 0.81, *p* = 0.41; BF_+0_ = 0.23]. However, an analysis with the early phase of the reconditioning (middle panel) revealed that they showed a significantly smaller CR to the CS+_extinction_ relative to CS- [*t*(40) = -3.11, *p* = 0.003; BF_+0_ = 10.34] while there was no significant difference between responses to CS+_active_ and CS- (i.e., CR) [*t*(40) = -0-25, *p* = 0.801; BF_+0_ = 0.17]. This suggests that the EXT training may better prevent reconditioning in the early phase. Yet, such effect was transient as participants did not show any CR to either the CS+_extinction_ or to the CS+_active_ in the late phase of reconditioning (right panel) [CS+_extinction_; *t*(40) = 1.04, *p* = 0.305; BF_+0_ = 0.7; CS+_active_; *t*(40) = 1.45, *p* = 0.15; BF_+0_ = 0.44]. The CR did not differ between the CS+s in the late phase of reconditioning [*t*(40) = 1.108, *p* = 0.27; BF_+0_ = 0.29]. Data are represented as mean ± between subjects SEM. *\*\*p < 0.01*

**Figure S6.**
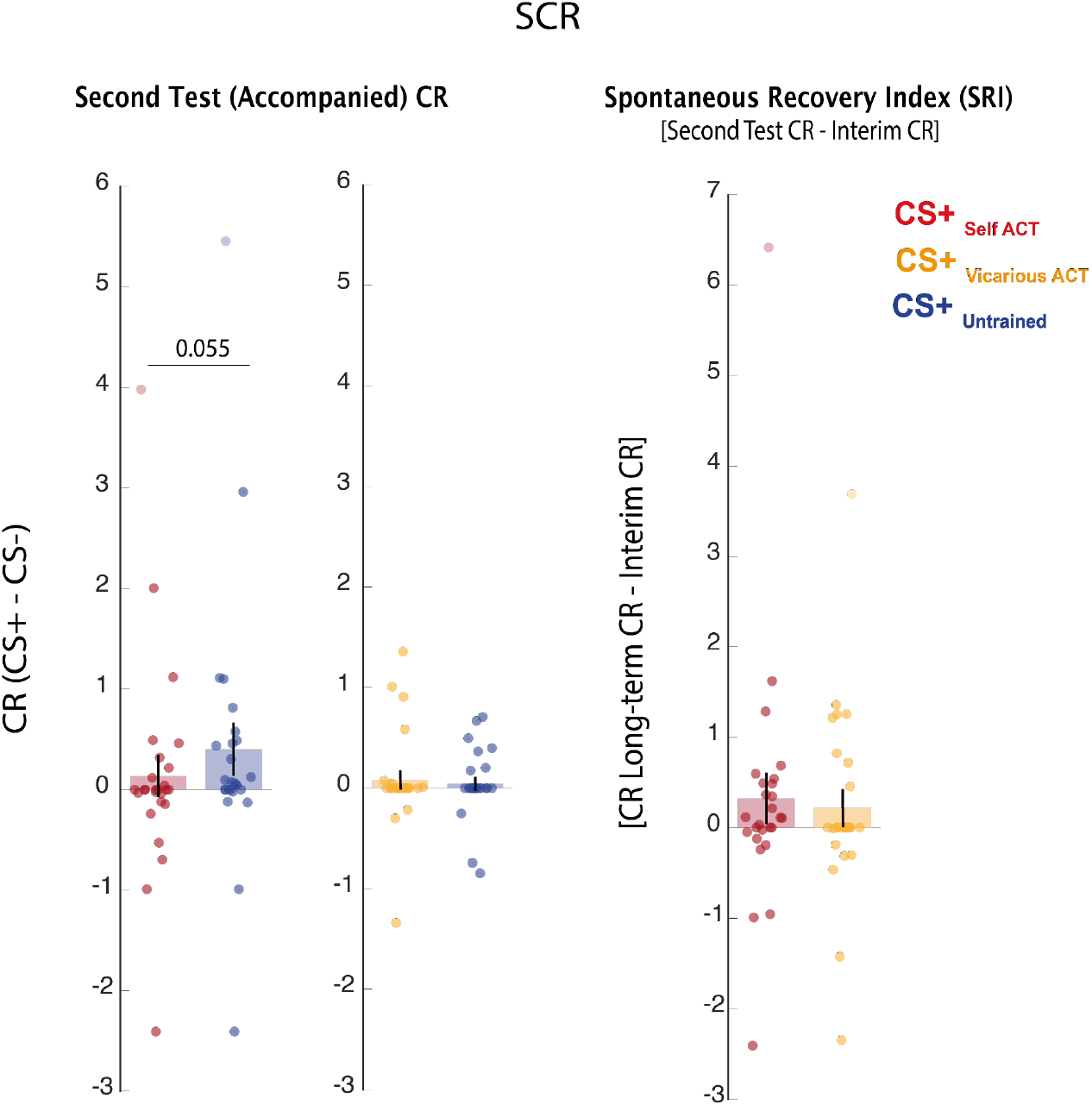
SCR results for the Second Test in Group 2. Self and Vicarious training on conditioned SCR (left and middle) and spontaneous recovery index (SRI) (right) in the second test in Group 2. In the second test, participants were accompanied by the third avatar from the training sessions. Although the results were qualitatively similar to the spontaneous recovery test, we did not find a significant interaction between training type and CS+ type [F(1,50) = 2.27 *p* = 0.13]. Data are represented as mean ± between subjects SEM.

**Figure S7.**
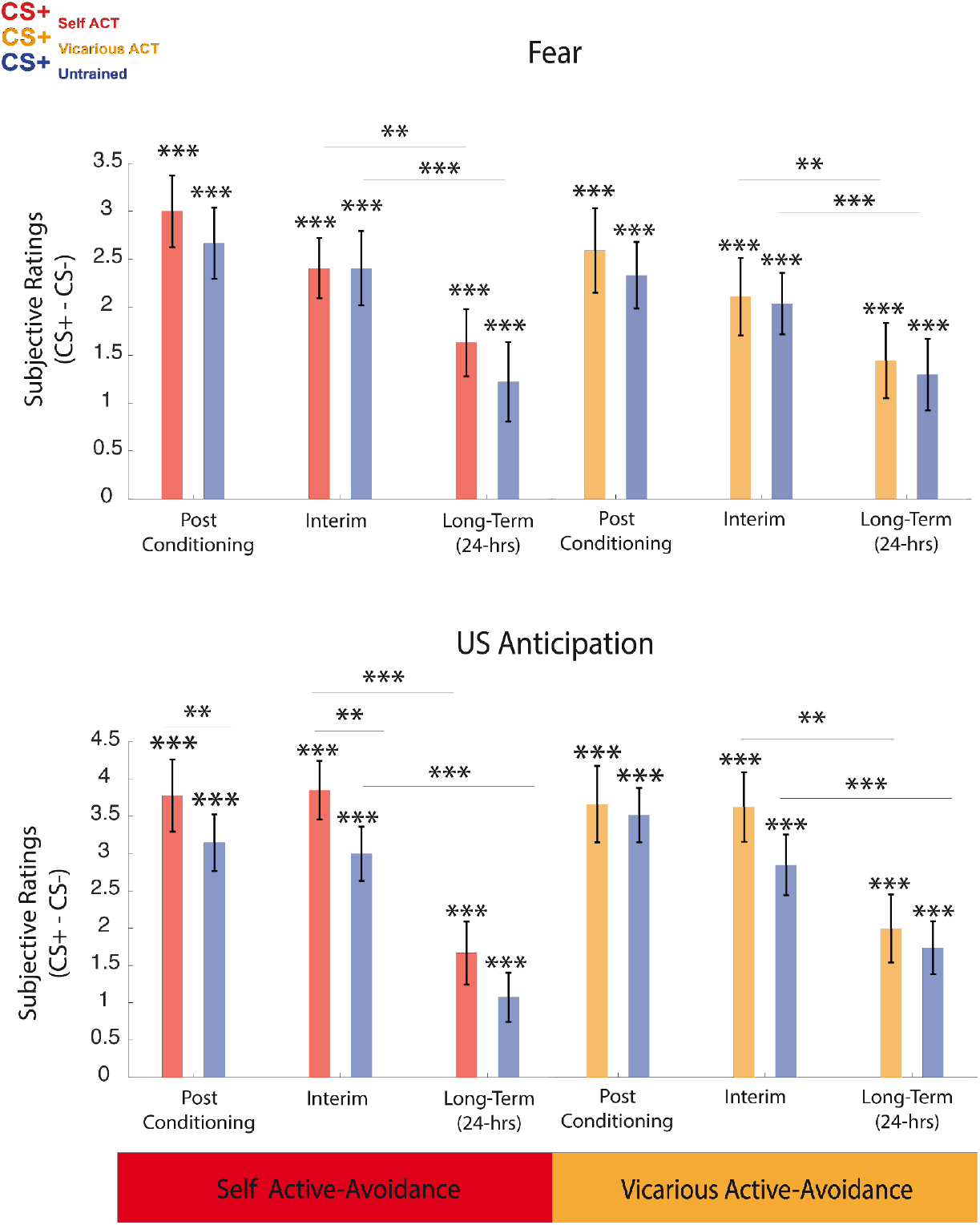
Comparison of fear alleviation effects between Self and Vicarious ACT training on subjective ratings. Interim and long-term fear reduction effects of Group 2 were also assessed with subjective ratings. Both Self- and Vicarious-groups reported higher feelings of fear as well as higher anticipation of violence toward both CS+_active_ and CS+_untrained_ relative to CS- after threat conditioning on day 1, indicating successful threat conditioning. However, Self-group reported higher anticipation of violence toward CS+_active_ than CS+_untrained_. Next, we compared the interim and long-term fear alleviating effects between the two types of ACT training on day 1 and day 2, respectively. In both groups, the fear ratings to the CS+_active_ and CS+_untrained_ on day 1 (post-training) and day 2 (post-test) were similar, suggesting a comparable interim and long-term fear reduction effect of both ACT training relative to no-training. Accordingly, the anticipation of violence toward the CS+_active_ and CS+_untrained_ on day 1 (post-training) and day 2 (post-test) were also similar, but Self-group showed higher anticipation of violence toward CS+_active_ than CS+_untrained_ as in the post-threat conditioning. Similarly, both fear ratings and anticipation of violence decreased from day 2 (post-training) to day 3 (post-test) in both groups regardless of the presence of the ACT training. These results suggest that the ACT training effects on subjective ratings may generalise to CS+ receiving no other training. Data are represented as mean ± between subjects SEM. *\* correspond to classical paired t-tests. **p < 0.01 ***p < 0.001*.

**Movie S1.**

CS+ (US)

**Movie S2.**

ACT training

**Movie S3.**

CS+ (no US) or EXT training

